# Ablation of SUN2-containing LINC complexes drives cardiac hypertrophy without interstitial fibrosis

**DOI:** 10.1101/612150

**Authors:** Rachel M. Stewart, Elisa C. Rodriguez, Megan C. King

## Abstract

The cardiomyocyte cytoskeleton, including the sarcomeric contractile apparatus, forms a cohesive network with cellular adhesions at the plasma membrane and nuclear-cytoskeletal linkages (LINC complexes) at the nuclear envelope. Human cardiomyopathies are genetically linked to the LINC complex and A-type lamins, but an full understanding of disease etiology in these patients is lacking. Here we show SUN2-null mice display cardiac hypertrophy coincident with enhanced AKT/MAPK signaling, as has been described previously for mice lacking A-type lamins. Surprisingly, in contrast to lamin A/C-null mice, SUN2-null mice fail to show coincident fibrosis or upregulation of pathological hypertrophy markers. Thus, cardiac hypertrophy is uncoupled from pro-fibrotic signaling in this mouse model, which we tie to a requirement for the LINC complex in productive TGFβ signaling. In the absence of SUN2, we detect elevated levels of the integral inner nuclear membrane protein MAN1, an established negative regulator of TGFβ signaling, at the nuclear envelope. We suggest that A-type lamins and SUN2 play antagonistic roles in the modulation of pro-fibrotic signaling through opposite effects on MAN1 levels at the nuclear lamina, suggesting a new perspective on disease etiology.

## Introduction

The mammalian myocardium is comprised of cardiomyocytes, which contain sarcomeres, the basic structural unit of muscle. Sarcomeres form a cohesive, tissue-scale network between cell-cell adhesions at the intercalated disc (ICD) and cell-extracellular matrix adhesions at costameres in these cells. Embedded into the contractile network of cardiomyocytes is the nucleus, which is mechanically integrated into the cytoskeleton through nuclear envelope-spanning LINC (Linker of Nucleoskeleton and Cytoskeleton) complexes, which consist of SUN domain proteins in the inner nuclear membrane and KASH domain proteins, Nesprins or SYNEs in mammals, in the outer nuclear membrane (Chang *et al.*, 2015). These complexes interface directly and indirectly with all components of the cytoskeleton on their cytoplasmic face, and interact with the nuclear lamina on their nucleoplasmic face (Chang *et al.*, 2015). While it is well-established that the formation and maintenance of cellular adhesions is dependent on proper cytoskeletal function (Reviewed in (Sequeira *et al.*, 2014)), our previous work (and that of others) implicates LINC complexes and the nuclear lamina as unexpected regulators of cellular adhesions at the plasma membrane (Mounkes *et al.*, 2005; Frock *et al.*, 2012; Stewart *et al.*, 2015). In addition to their structural roles, adhesions and components of the sarcomere are also mechanoresponsive and play an important role in the biochemical signaling processes that promote cardiac function, such as the normal hypertrophic growth that occurs during cardiac development (Reviewed in (McCain and Parker, 2011; Maillet *et al.*, 2013)). Thus, it is not surprising that defects in adhesion are strongly implicated in human cardiomyopathies (Sheikh *et al.*, 2009; Delmar and McKenna, 2010; Israeli-Rosenberg *et al.*, 2014).

The LINC complex and other components of the nuclear lamina, such as emerin and A-type lamins, are also genetically linked to various human myopathies, including dilated cardiomyopathy, arrhythmogenic cardiomyopathy, and syndromes with cardiac involvement such as Emery-Dreifuss muscular dystrophy (EDMD) (Ostlund *et al.*, 2001; Stroud *et al.*, 2014). Strikingly, a large proportion of individuals with dilated cardiomyopathies exhibit lamin A/C point mutations, preceded in prevalence only by mutations in titin (Arbustini *et al.*, 2002; Cahill *et al.*, 2013). Point mutations in *Sun2*, one of the most widely expressed genes encoding a SUN domain in mammals, have been found to act as genetic modifiers of cardiomyopathy, for example, exacerbating the severity of the disease in patients with mutations in the sarcomere component myosin binding protein C or other components of the nuclear lamina (Meinke *et al.*, 2014). These point mutations in *Sun2* reside in either the lamin A-binding region (M50T) or in the coiled-coil region, required for the trimerization of LINC complexes and Nesprin engagement (V378T) (Sosa *et al.*, 2012; Meinke *et al.*, 2014). Further, SUN2 V378T and other point mutations in the coiled-coil region have been found in patients with EDMD or cardiomyopathy symptoms (Meinke *et al.*, 2014). Despite the strong association between disruption of the LINC complex or nuclear lamina and human cardiac disease, a mechanistic understanding of the origin of these genetic diseases remains elusive. One prevailing model focuses on characteristic changes in nuclear morphology that accompany disruption of nuclear lamina components (Mounkes *et al.*, 2005; Gupta *et al.*, 2010; Zwerger *et al.*, 2013), suggesting that nuclear fragility is the driving force behind disease etiology (Reviewed in (Davidson and Lammerding, 2014)).

An additional (or alternative) mechanistic link has recently come to light from studies employing mouse models of lamin depletion (or a cardiomyopathy-linked lamin A/C mutation - H222P). Here, several studies uncovered altered MAP kinase and AKT signaling as a driver of cardiac dysfunction (Muchir *et al.*, 2007; Wu *et al.*, 2011; Choi *et al.*, 2012; Muchir *et al.*, 2012b); reversing this biochemical cascade by treatment with the TOR inhibitor rapamycin improved heart function (Choi *et al.*, 2012; Liao *et al.*, 2016). However, the mechanism(s) by which mutations in lamin A (or loss of lamin A) alters the AKT and TOR pathways remains unknown. Importantly, both the AKT and the pro-fibrotic TGFβ pathways that drive cardiac hypertrophy and interstitial fibrosis are mechanoresponsive, largely through pathways sensitive to inputs from cellular adhesions and/or the state of the actomyosin cytoskeleton (Iijima *et al.*, 2002; Balasubramanian and Kuppuswamy, 2003; Gomez *et al.*, 2010; Young *et al.*, 2014; Hinz, 2015; O’Connor *et al.*, 2015; Varney *et al.*, 2016). Models of LINC complex ablation provide an opportunity to test whether communication of mechanical signals from the cytoskeleton to the nuclear interior (Lammerding *et al.*, 2004; Lee *et al.*, 2007; Hale *et al.*, 2008; Stewart-Hutchinson *et al.*, 2008; Luxton *et al.*, 2010a; Folker *et al.*, 2011; Khatau *et al.*, 2012; Long *et al.*, 2013; Myat *et al.*, 2015; Stewart *et al.*, 2015) contributes to myocardium function, although to date this avenue of investigation has been largely unexplored.

Although suppressing AKT and MAPK signaling improves cardiac performance in lamin A/C-null mice, this intervention does not counteract the coincident fibrosis (Muchir *et al.*, 2007; Wu *et al.*, 2011; Choi *et al.*, 2012; Muchir *et al.*, 2012b). Moreover, fibrosis is a hallmark of the dysfunction characteristic of other laminopathic syndromes, including progeria and lipodystrophy (Olive *et al.*, 2010; Le Dour *et al.*, 2017). While the underlying mechanisms are poorly understood, it is interesting to note that MAN1, an integral inner nuclear membrane (INM) protein that requires A-type lamins to be retained at the INM (Ostlund *et al.*, 2006), is a clear negative regulator of TGFβ signaling across a broad range of organisms (Raju *et al.*, 2003; Lin *et al.*, 2005; Ishimura *et al.*, 2006) In addition to binding directly to R-SMADs, MAN1 has also been suggested to regulate SMAD signaling in a tissue stiffness-dependent manner (Chambers *et al.*, 2018).

Here, to address how LINC complexes contribute to heart function, we have examined the consequences of disrupting SUN2-containing LINC complexes in the murine myocardium, and find that this genetic perturbation induces cardiac hypertrophy. *Sun2−/−* mice display elevated AKT-mTOR and MAPK signaling in the myocardium, which we tie to increased integrin engagement at costameres. Surprisingly, these mice fail to induce expression of classic hypertrophy-associated genes, have a normal lifespan, lack fibrosis, and demonstrate down-regulation or unaltered levels of TGFβ target genes despite elevated levels of a transducer of this pathway, nuclear phospho-SMAD2. While lamin A/C is required for MAN1 targeting, we find that SUN2-null mice instead display elevated retention of MAN1 at the nuclear lamina. Taken together, these results suggest that A-type lamins and the LINC complex act in concert to regulate pro-hypertrophic signaling, but play antagonistic roles in driving fibrosis.

## Results

### Mice deficient for *Sun2* undergo cardiac hypertrophy

To assess the functional consequences of *Sun2* loss in the murine myocardium, we obtained a previously reported *Sun2−/−* whole body knockout mouse model (Lei *et al.*, 2009). In wildtype (WT) left ventricular cardiomyocytes, SUN2 is localized to the nuclear envelope and is absent from *Sun2−/−* tissue (Supplemental Figure 1A); SUN1 expression is not substantially different in the hearts of *Sun2−/−* mice compared to WT (Supplemental Figure 1B). While we did not observe increases in spontaneous cardiac-associated deaths in aged mice (>1 year), gross histology of hearts cut at the mid-ventricular level revealed enlargement of *Sun2−/−* hearts compared to WT hearts at over one year of age (Figure 1A). These findings were recapitulated at the cellular level, as we observed significant enlargement of individual cardiomyocytes in the papillary muscle of *Sun2−/−* mice (Figure 1B, C). These results suggest that *Sun2−/−* mice exhibit age-related cardiac hypertrophy at both the cellular and tissue levels.

**Figure 1.**
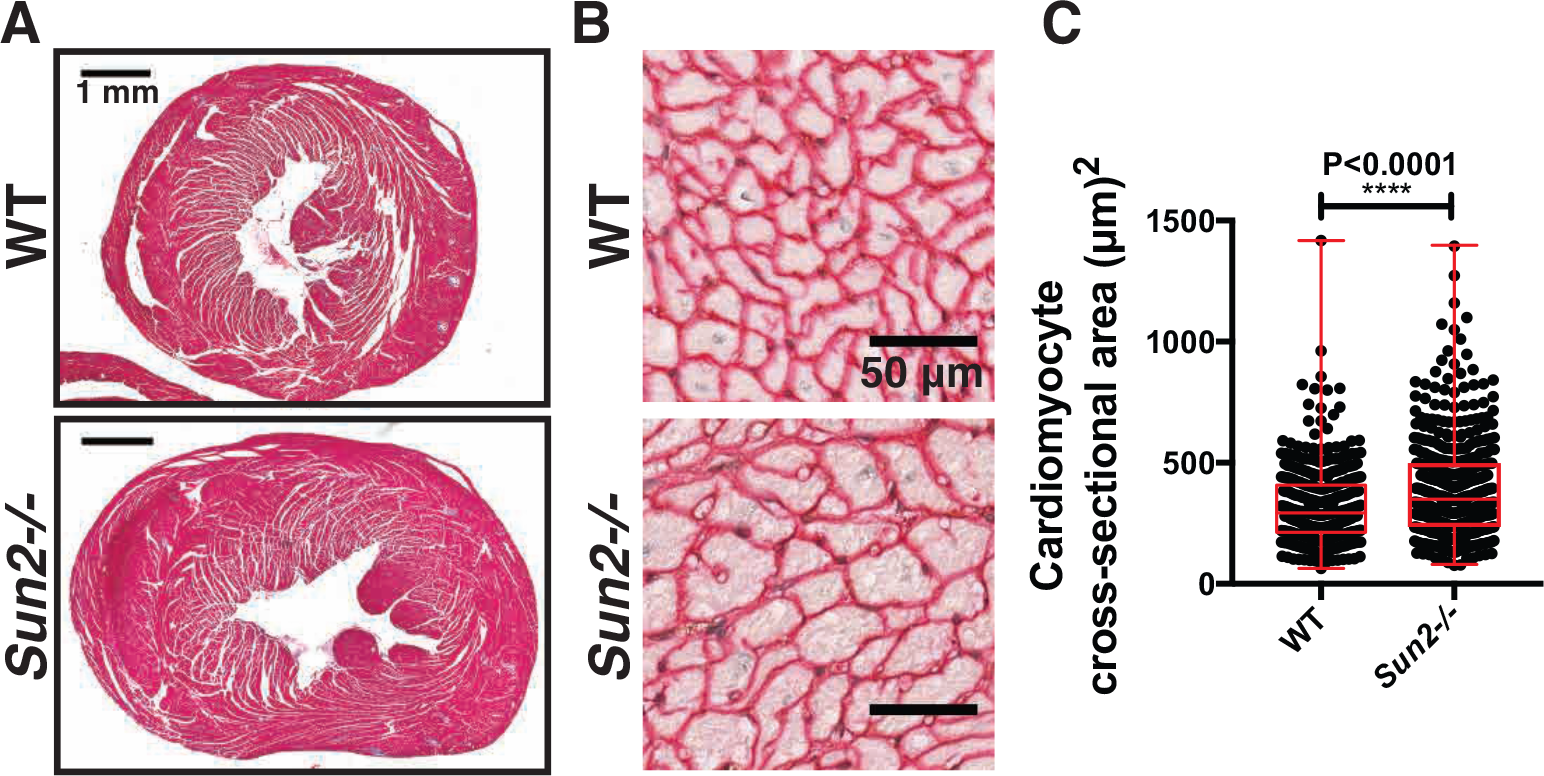
*Sun2−/−* murine hearts exhibit hypertrophy. (A) Paraffin-embedded hearts isolated from 13-month-old WT and *Sun2−/−* mice were stained with Masson’s trichrome. Representative images show enlargement of the *Sun2−/−* heart compared to WT; images of additional hearts are displayed in Supplemental Figure 1C. (B) Paraffin-embedded hearts from WT and *Sun2−/−* mice were stained with antibodies against laminin to reveal cardiomyocyte outlines. Cardiomyocytes from left ventricular papillary muscle are shown in cross-section. Note the enlargement of *Sun2−/−* cells compared to WT. (C) Quantification of left ventricular papillary muscle cardiomyocyte cross-sectional area, showing a greater population of enlarged cells in *Sun2−/−* compared to WT heart. n = ≥86 cells (86 – 198 cells) for each of 3 mice per genotype. Error bars indicate SDs. Statistical significance determined by unpaired, two-tailed *t* test.

### *Sun2−/−* mice exhibit altered sarcomere structure and adhesion defects

Cardiac dysfunction is often tied to changes in sarcomere structure. In particular, myofibril disarray has been linked to sarcomere mutations, many of which drive increased contractile function of the sarcomere at the cellular level (Michele *et al.*, 1999; Lowey, 2002; Moore *et al.*, 2012). Assessment of sarcomere organization in P50 mice by staining left ventricle (LV) cardiac tissue sections with phalloidin revealed the stereotypical banding of F-actin, corresponding to I-bands, in WT mice (Figure 2A). However, *Sun2−/−* tissue displayed actin bands of irregular width that did not align laterally between adjacent myofibrils, as well as regions with extensive actin disorganization (Figure 2A arrowheads). At higher magnification by transmission electron microscopy (TEM), we find that while many regions of the *Sun2−/−* tissue still exhibited grossly intact myofibril structure (Figure 2B), these regions displayed misaligned and wavy Z-bands (Figure 2B, red lines) and M-bands (Figure 2B, arrowheads), loss of clearly defined I-bands (Figure 2B, arrows) and H-zones (Figure 2B, arrowheads), and reduced sarcomere length (Figure 2C). Focal regions of severe myofibril disarray with complete loss of sarcomere structure were also observed in the *Sun2−/−* LV (Figure 2B, lower panels).

**Figure 2.**
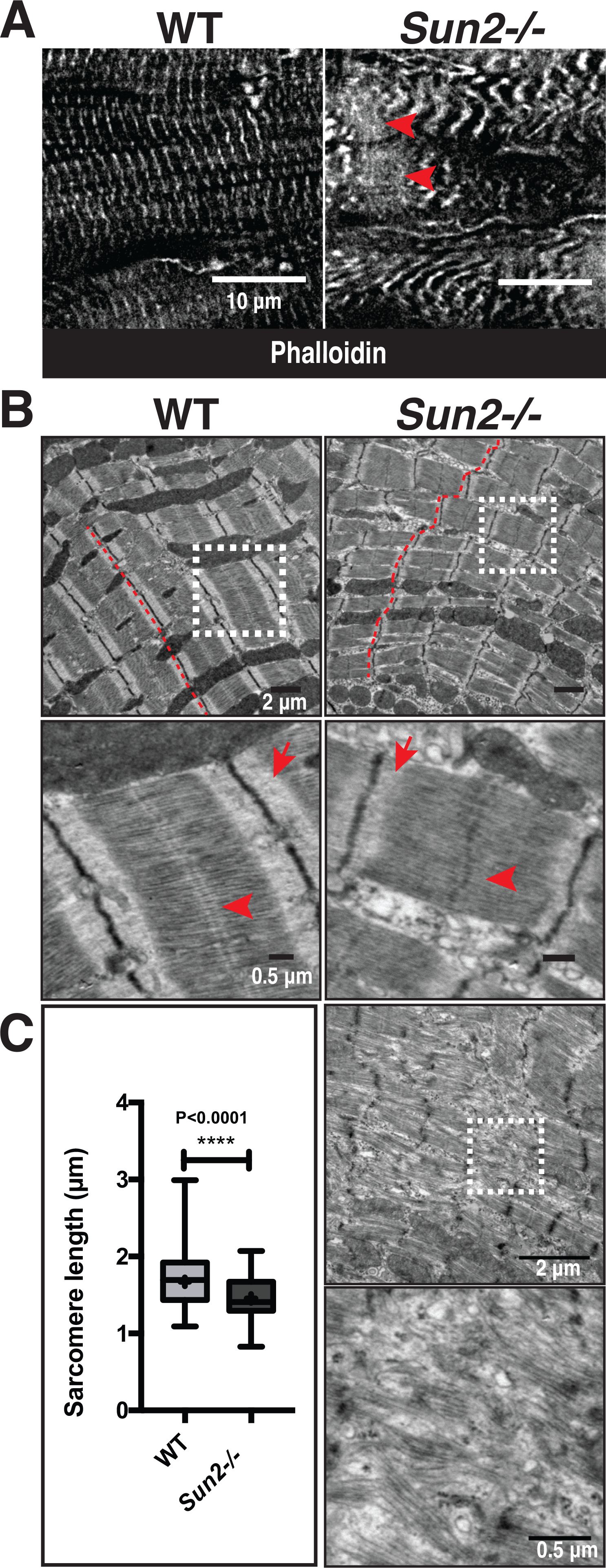
Sarcomere organization is perturbed in P50 *Sun2−/−* mice. (A) Frozen P50 WT and *Sun2−/−* cardiac left ventricle tissue was sectioned and stained with phalloidin to label F-actin in I-bands. While I-bands are evenly registered in WT tissue, the regularity of this pattern is lost in the *Sun2−/−*. Note the variability in width of I-bands, loss of alignment and increased wavy appearance of I-bands, and focal loss of sarcomere structure (arrowheads). (B) Transmission electron micrographs (TEM) of P50 WT and *Sun2−/−* left ventricle tissue. Classical sarcomere structure can be observed in WT mice, while *Sun2−/−* tissue exhibits misaligned and wavy Z-bands (compare red lines in WT versus *Sun2−/−*) and loss of clearly defined I-bands (arrows). Focal regions of complete sarcomere disruption are also present in the *Sun2-/* tissue (bottom two panels). (C) Sarcomere length, as measured from Z-band-to-Z-band, is reduced in P50 *Sun2−/−* tissue. n = 3 mice for each genotype. Statistical significance determined by unpaired, two-tailed *t* test.

Our previous work showing a requirement for SUN2 LINC complexes in intercellular adhesion organization and function (Stewart *et al.*, 2015) inspired us to examine the structure of intercalated discs, the intercellular adhesions that connect adjacent cardiomyocytes. TEM analysis revealed discontinuous jagged intercalated discs with lacunae where sarcomeric cytoskeletal filaments failed to properly interface with intercalated disc components in P50 *Sun2−/−* tissue (Supplemental Figure 2A, arrows). Further, we observed reduced electron density along the intercalated disc membranes, suggesting that there was reduced adhesion protein accumulation at these sites (Supplemental Figure 2A, arrowheads).

Immunofluorescence staining of intercalated disc components desmoplakin I/II, corresponding to desmosomes, and β-catenin, corresponding to adherens junctions, revealed discrete bands that ran perpendicular to the cardiomyocyte lateral membranes in WT tissue (Supplemental Figure 2B,C). However, we frequently observed expansion of these normally discrete bands into several thinner, separated bands in the *Sun2−/−* tissue (Supplemental Figure 2B,C, arrowheads), which mirrored the discontinuous appearance of the intercalated discs at the level of TEM (Supplemental Figure 2A). These results suggest that intercellular adhesion is structurally perturbed in the absence of SUN2 LINC complexes.

### Sarcomere disorganization precedes changes in nuclear morphology in *Sun2−/−* mice

By P50 we observed extensive changes in sarcomere organization, and cardiomyocyte adhesion in *Sun2−/−* mice. To address the likely foundational defect that drives these phenotypes, we next investigated early time points in mouse development. Further, as we observed altered nuclear morphology in P50 *Sun2−/−* cardiac tissue (Figure 3A,C), which has been suggested to drive disease in laminopathies and nuclear envelopathies (Mounkes *et al.*, 2005; Gupta *et al.*, 2010; Zwerger *et al.*, 2013), we also wished to investigate if changes in the myocardium ultrastructure preceded or followed changes in nuclear shape. Morphological changes in nuclear shape were not present at P4 (Figure 3B,C), suggesting that defects in nuclear morphology arise only after mature cardiac beating begins. We did observe that P4 *Sun2−/−* nuclei exhibited a somewhat rounder morphology than the more elongated WT nuclei (Figure 3B); however, we would argue that this observation likely reflects that LINC complex-dependent tension is normally exerted by the cytoskeleton on the nucleus in WT cardiomyocytes, a phenomenon that may be lost upon ablation of LINC complexes (Zwerger *et al.*, 2013; Hatch and Hetzer, 2016).

**Figure 3.**
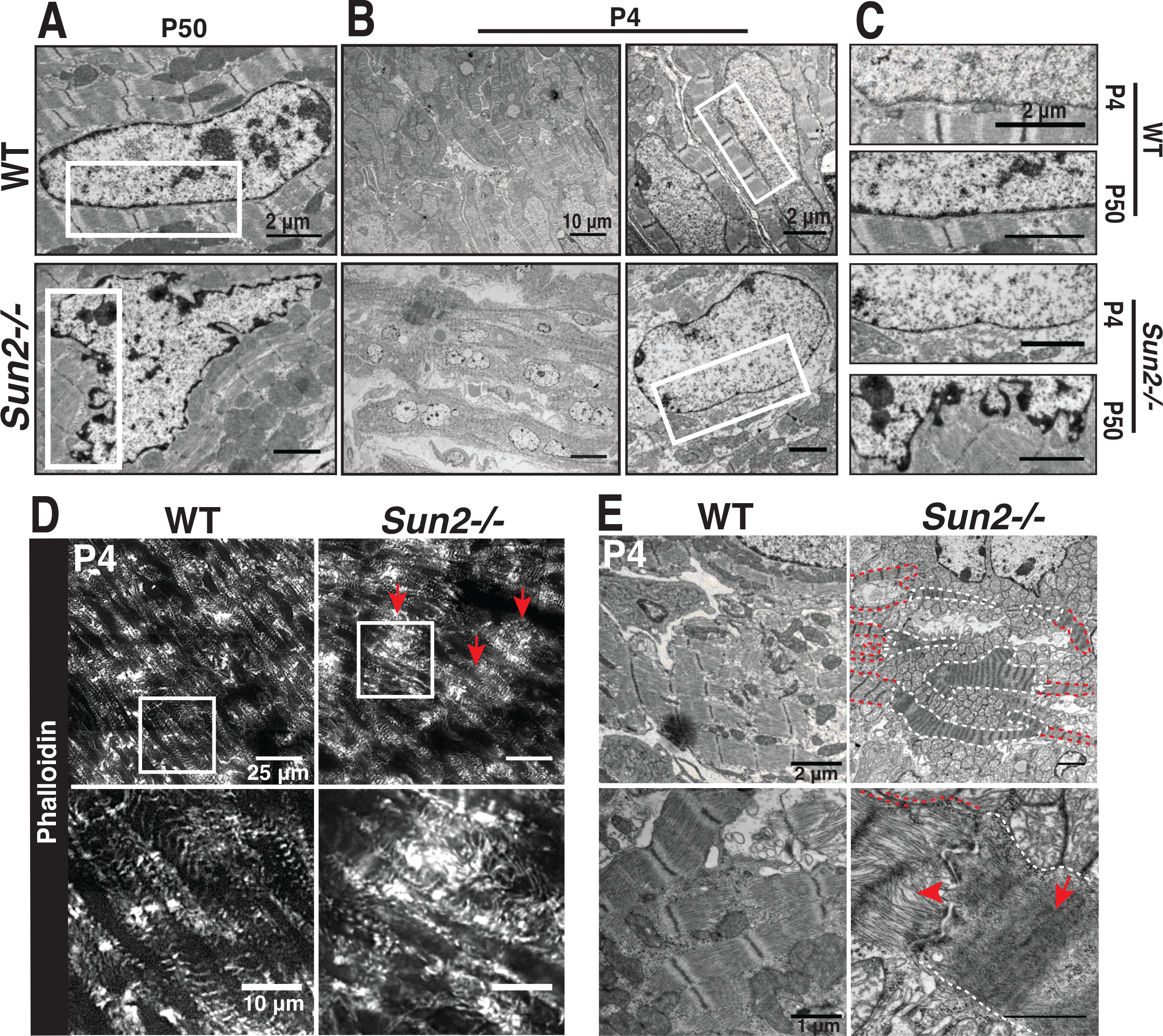
Nuclear morphology is normal at P4 but sarcomere organization is disrupted in *Sun2−/−* mice. (A-C) TEM of WT and *Sun2−/−* left ventricle tissue at P50 and P4. (A) At P50, nuclei in *Sun2−/−* tissue exhibit severe distortions of the nuclear envelope. (B) At P4, *Sun2−/−* nuclear envelope morphology is comparable to the morphology in WT tissue. Left panels display lower magnification fields, illustrating the prevalence of morphologically-similar nuclei in WT and *Sun2−/−* tissue. However, WT nuclei are more elongated than *Sun2−/−* nuclei. Right panels display higher magnification of single nuclei, highlighting the similarity in nuclear envelope structure in the two genotypes. (C) Insets from white boxed regions in (A) and (B), illustrating the normal nuclear morphology at P4 and disruption by P50 in *Sun2−/−* tissue. (D) Frozen P4 WT and *Sun2−/−* cardiac left ventricle tissue was sectioned and stained with phalloidin. While nascent myofibrils in WT tissue display irregularity in I-band width and alignment, focal regions of *Sun2−/−* tissue exhibit severe sarcomere disruption (arrows). (E) TEM of P4 WT and *Sun2−/−* left ventricle tissue. Sarcomere structure is disrupted in *Sun2−/−* tissue, with cytoskeletal disorganization (arrowhead) and hypercontraction (arrow) visible in adjacent cells. Hypercontractile cells are outlined in white, while adjacent cells with more subtly disrupted sarcomere organization are outlined in red.

Given that the nuclei appeared intact in the *Sun2−/−* LV at P4, we next asked if the P50 sarcomere defects (Figure 2) manifest before or after this stage in development; in addition, the ICDs are not yet mature at P4 (Hirschy *et al.*, 2006; Wang *et al.*, 2012), further allowing us to examine if the sarcomere changes occurred prior to ICD formation. As many structures in the murine heart, including myofibrils and adhesions, continue to develop postnatally (Hirschy *et al.*, 2006; Wang *et al.*, 2012), phalloidin staining of P4 WT cardiac tissue revealed a less regular pattern of actin-rich I-bands than was observed at P50 (Figure 3D versus Figure 2A). However, at this earlier age we again observed extensive disruptions in myofibril organization in *Sun2−/−* tissue compared to WT tissue (Figure 3D, arrows). These defects were further clarified following TEM of P4 LV myocardium, where *Sun2−/−* tissue featured ragged sarcomeres with a loss of clearly discernible I-bands (Figure 3E, arrowhead) and irregular Z-band spacing (Figure 3E, arrow). This indicates that sarcomere disarray is present during early postnatal cardiac development in mice lacking *Sun2*, prior to changes in nuclear morphology and ICD maturation.

We identified several regions in the P4 *Sun2−/−* tissue with extensive, striking reductions in the spacing between Z-bands, a phenotype previously linked to hypercontractility of the sarcomere (Lauritzen *et al.*, 2009) (Figure 3E, arrow). It appears that this effect was cell autonomous, as individual cells with this hallmark of hypercontractility were located only sporadically through the examined tissue, while adjacent cells displayed more subtle changes in sarcomere structure (Figure 3E, compare hypercontractile cells outlined in white to adjacent cells outlined in red). In contrast to the P50 time point, we did not observe noticeable differences in intercellular adhesion in *Sun2−/−* mice during the two-week postnatal period when intercalated disc maturation occurs (Hirschy *et al.*, 2006; Wang *et al.*, 2012) (Supplemental Figure 2D). This suggests that the sarcomere disarray in *Sun2−/−* mice precedes the alterations in intercellular adhesion, and that the intercalated disc defects observed by P50 may be a consequence of altered contractility and/or the sarcomere defects in these mice.

### *Sun2−/−* mice exhibit increased integrin engagement and AKT/MAPK signaling

Given the established roles that costameres, the sites of cell-extracellular matrix adhesion, play in cardiac development, sarcomere organization, and integrin signaling upstream of the pro-hypertrophic AKT, mTOR, and MAPK pathways and their mechanoresponsive nature (MacKenna *et al.*, 1998; Franchini *et al.*, 2000; Laser *et al.*, 2000; Balasubramanian and Kuppuswamy, 2003; Brancaccio *et al.*, 2006; Dowling *et al.*, 2008; Konieczny *et al.*, 2008; McCain *et al.*, 2012; Israeli-Rosenberg *et al.*, 2014; Wilsbacher and Coughlin, 2015), we next examined costameres for functional changes in *Sun2−/−* mice. We employed a well-established antibody (9EG7) that recognizes a ligand-bound epitope of β1-integrin, thought to correspond to the active, engaged form of the protein (Lenter *et al.*, 1993; Bazzoni *et al.*, 1995; Su *et al.*, 2016). While levels of total β1-integrin were similar in WT and *Sun2−/−* P50 tissue, the levels of active 9EG7 β1-integrin appeared substantially higher in the *Sun2−/−* tissue (Figure 4A, see additional examples in Supplemental Figure 2E). We did not observe an increase in active β1-integrin levels at P13 (Supplemental Figure 2F) or in 13 month-old mice (Supplemental Figure 2G). These results suggest that integrin function at the costamere may be altered in the absence of SUN2 LINC complexes at a specific stage in murine cardiac development.

**Figure 4.**
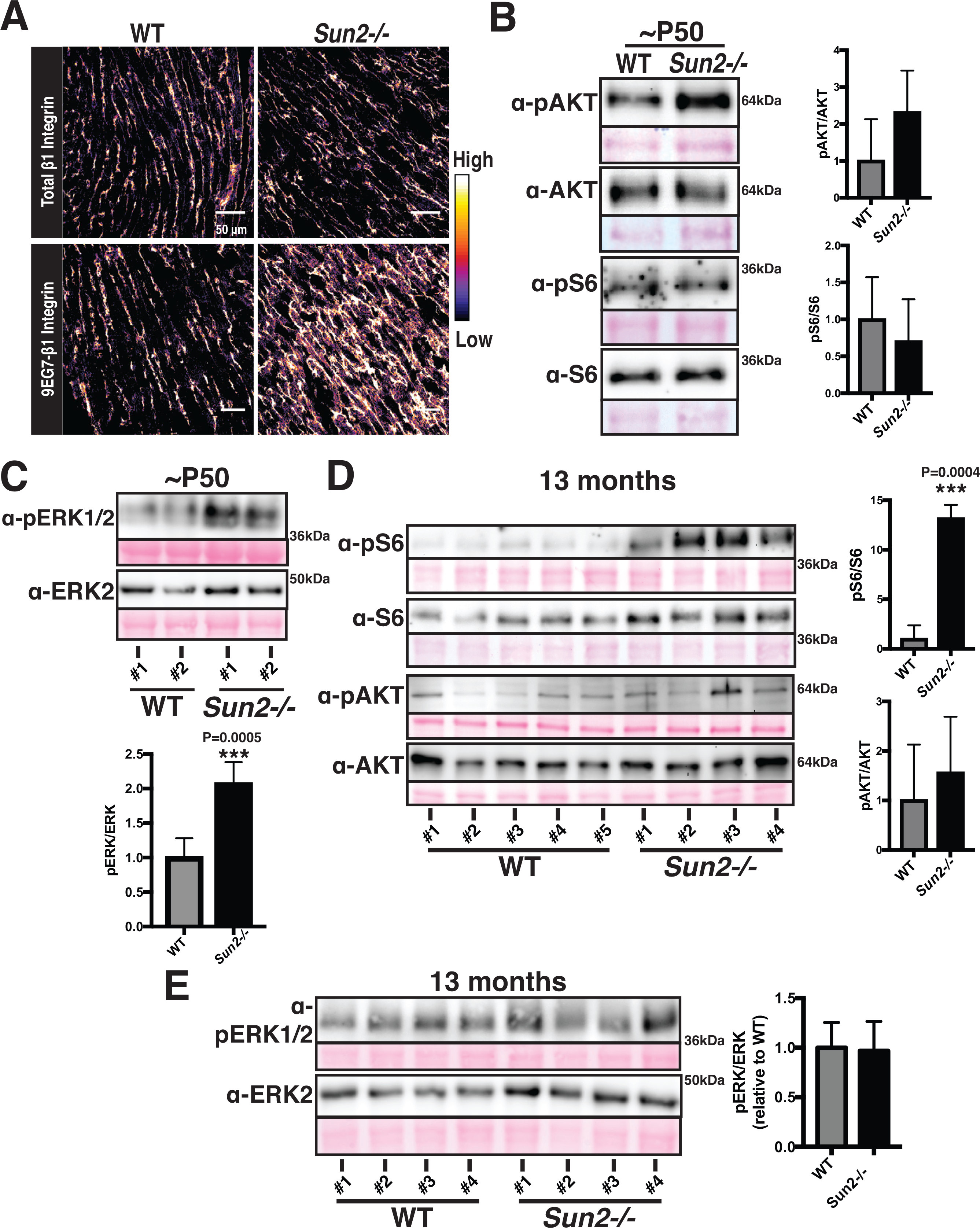
*Sun2−/−* hearts display increased integrin engagement and AKT/MAPK signaling. (A) Frozen P50 WT and *Sun2−/−* cardiac left ventricle tissue was sectioned and stained with antibodies against total β1-integrin (upper panels) or against the ligand-bound, active β1-integrin (9EG7) conformational epitope (lower panels). Images were pseudo-colored to depict the relative fluorescence intensity of the β1-integrin signal, with lighter colors indicative of higher intensity and darker colors indicative of lower intensity. Note the increased intensity of active β1-integrin in the *Sun2−/−* tissue. Images are representative of results for 3 mice per genotype (also see Supplemental Figure 2E-G). (B-E) Representative immunoblots of ventricular lysate from 3 WT and 3 *Sun2−/−* mice at P50 or 5 WT and 4 *Sun2−/−* mice at 13 months. Ponceau staining of total protein reveals even loading of samples. Plots to the right represent quantitative analysis from additional mice, represented as the mean ± SD for >3 WT or *Sun2−/−* mice. Full blots, used for quantifications, are shown in Supplemental Figure 3. For quantifications, data are shown as a ratio of *Sun2−/−* to WT for the phosphorylated protein and total protein, expressed as a ratio. (B) Lysates from P50 mice were subjected to SDS-PAGE and immunoblotting with antibodies against phosphorylated AKT (pAKT), AKT, phosphorylated S6 (pS6), or S6, revealing elevated levels of pAKT in *Sun2−/−* tissue. Data are represented as the mean ± SD for 3 WT or *Sun2−/−* mice. (C) Lysates from P50 mice were subjected to SDS-PAGE and immunoblotting with antibodies against pERK1/2 and ERK2, revealing elevated levels of pERK1/2 in *Sun2−/−* tissue. Data are represented as the mean ± SD for 3 WT or *Sun2−/−* mice. (D) Lysates from 13 month-old mice were subjected to SDS-PAGE and immunoblotting with antibodies against phosphorylated S6 (pS6), S6, phosphorylated AKT (pAKT), or AKT, revealing elevated levels of pS6 but similar levels of pAKT in *Sun2−/−* tissue versus WT tissue. Data are represented as the mean ± SD for 5 WT or 4 *Sun2−/−* mice. (E) Lysates from 13 month-old mice were subjected to SDS-PAGE and immunoblotting with antibodies against pERK1/2 and ERK2, revealing similar levels of pERK1/2 in *Sun2−/−* and WT tissues. Data are represented as the mean ± SD for 3 WT or *Sun2−/−* mice.

Hypertrophy can be driven by integrin signaling, including through the AKT and MAPK pathways (MacKenna *et al.*, 1998; Franchini *et al.*, 2000; Laser *et al.*, 2000; Kim *et al.*, 2003; Rota *et al.*, 2005; Cittadini *et al.*, 2006; Shiojima and Walsh, 2006; Catalucci *et al.*, 2009; Maillet *et al.*, 2013). Moreover, AKT activation has been linked to increased cardiac function and cardiomyocyte contractility (Kim *et al.*, 2003; Rota *et al.*, 2005; Cittadini *et al.*, 2006; Catalucci *et al.*, 2009). At the time point when we observe greater integrin engagement (P50), immunoblot analysis of WT and *Sun2−/−* myocardium revealed increased levels of phosphorylated AKT (Figure 4B) and phosphorylated ERK1/2 (Figure 4C) in *Sun2−/−* animals at P50. Further, in hypertrophic 13 month-old *Sun2−/−* mice we observed a dramatic increase in the level of phosphorylated ribosomal protein S6 (Figure 4D), a key target downstream of the AKT and mTOR pathways linked to the hypertrophic response (Shiojima and Walsh, 2006; Maillet *et al.*, 2013). At this 13 month time point, when *Sun2−/−* mice no longer display increased integrin engagement (Supplemental Figure 2G), we also failed to observe a significant increase in AKT phosphorylation (Figure 4D) or pERK1/2 levels (Figure 4E).

### Loss of SUN2 uncouples AKT- and MAPK-driven hypertrophy from cardiac fibrosis

In the vast majority of disease contexts, cardiac hypertrophy occurs coincident with interstitial fibrosis (Ho *et al.*, 2010), including in the laminopathic mouse models (Muchir *et al.*, 2007; Wu *et al.*, 2011; Choi *et al.*, 2012; Muchir *et al.*, 2012b). Importantly, the consequences of disrupting lamin A function are quite severe, leading to death even before maturity is reached in the case of the lamin A/C-null model. Given the myriad cytological defects apparent in the myocardium of *Sun2−/−* mice, the gains in AKT- and MAPK-signaling, and cardiac hypertrophy, we were surprised that we failed to observe early death in the SUN2-null model. Remarkably, we found no increase in interstitial fibrosis in 13 month-old *Sun2−/−* mice as assessed by Masson’s trichrome staining of collagen and quantification with a color-based pixel count algorithm (Figure 5A-B, see Methods). Moreover, pathological cardiac hypertrophy associated with fibrosis is typically accompanied by increased transcription of a number of fetal program genes involved in sarcomerogenesis, including in the lamin A/C H222P mouse model (Kim *et al.*, 2007; Taegtmeyer *et al.*, 2010). However, quantitative polymerase chain reaction (qPCR) revealed no significant increases in the expression of ANP, BNP, skeletal α-actin, or Serca2 in *Sun2−/−* mice compared to WT mice (Figure 5C), suggesting that loss of SUN2 drives hallmarks of physiological rather than pathological hypertrophy (Reviewed in (Maillet *et al.*, 2013)). We next assessed the levels of gene targets of TGFβ, which is a master regulator of pro-fibrotic signaling, at both P50 and at 13 months. At P50 we observed a consistent down-regulation of TGFβ-associated pro-fibrotic genes, including fibronectin (FN1) and the collagen genes COL1A1 and COL3A1 (Figure 5D). This does not appear to a reflect a genome-wide effect on the transcriptome, as a subset of TGFβ targets (FLNA) and non-targets (DES, which encodes Desmin) were unaltered (Figure 5D). At 13 months, levels of the FN1 and COL1A1 transcripts were similar between WT and *Sun2−/−* mice (Figure 5E), consistent with the lack of fibrosis indicated by trichome staining (Figure 5A-B), although we note that COL3A1 levels are significantly up-regulated in mice lacking SUN2.

**Figure 5.**
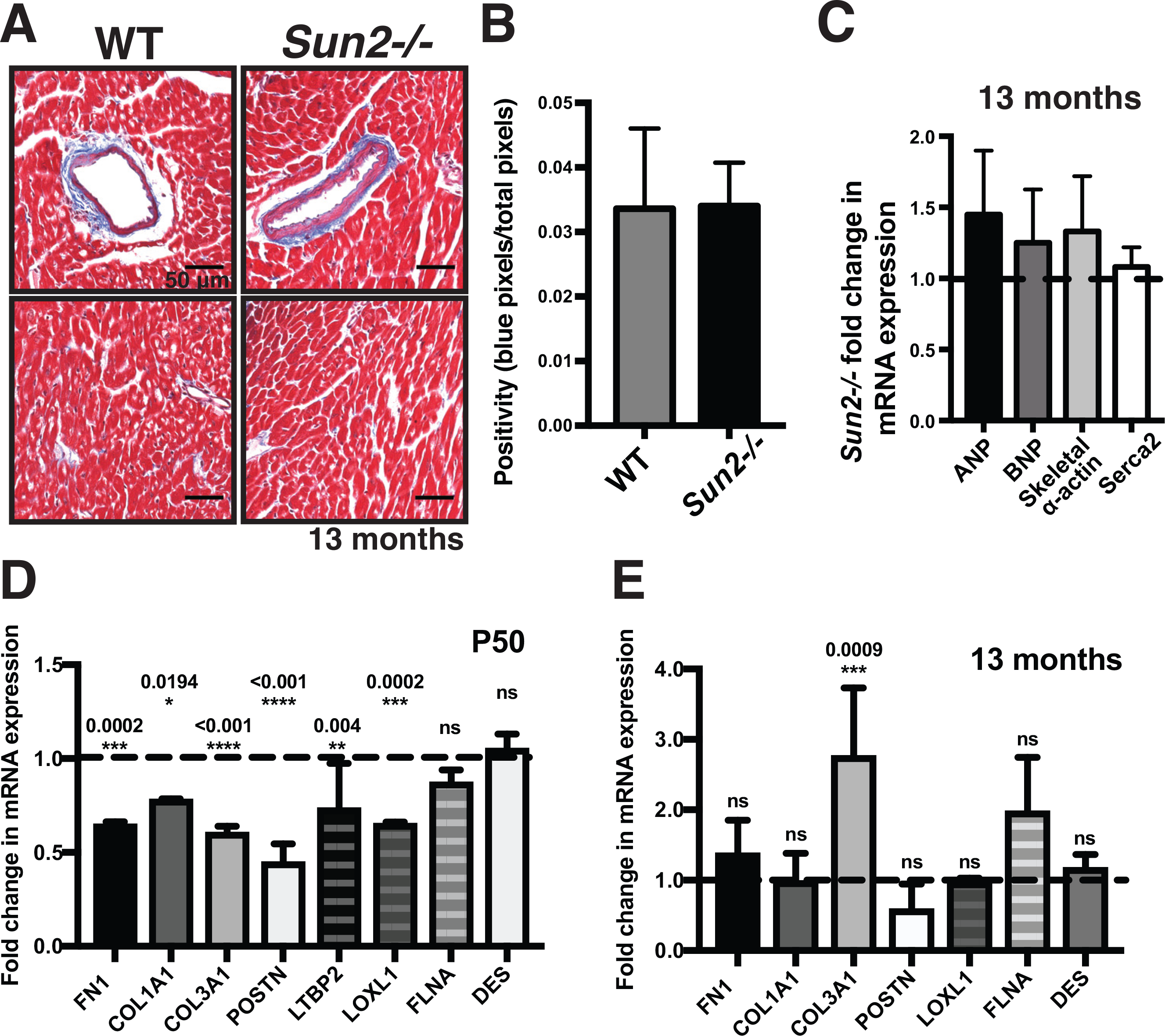
*Sun2−/−* hearts do not exhibit fibrosis and show reduced or unaltered expression of TGFβ target genes. (A) Images of trichrome-stained WT and *Sun2−/−* hearts are insets taken from Figure 1A. *Sun2−/−* tissue exhibits equivalent levels of collagen-positive staining (blue) as WT tissue surrounding blood vessels (top panels) and at interstitial (bottom panels) sites. (B) Quantification of collagen-staining positivity (the ratio of the number of blue pixels to total pixels, see Methods) in interstitial sites, showing equivalent levels of interstitial fibrosis in WT and *Sun2−/−* tissue. Unpaired, two-tailed *t* test failed to determine any difference as significant. (C) qPCR analysis of ANP, BNP, skeletal α-actin, and Serca2 mRNA levels from 13-month-old left ventricular total RNA. WT and *Sun2−/−* Ct values were normalized to GAPDH. Fold change of *Sun2−/−* mRNA levels was determined by calculating the 2^ΔΔCt^ after taking the mean of WT ΔCt values from 5 mice. n = 4 *Sun2−/−* mice. Lack of statistical significance as determined by unpaired, two-tailed *t* test. (D) qPCR analysis of FN1 (fibronectin), COL1A1 (collagen type I), COL3A1 (collage type III), POSTN (periostin), LTBP2 (latent TGFβ-binding protein 2), LOXL1 (lysyl oxidase homolog 1), and FLNA (filamin A) in P50 left ventricular total RNA, revealing similar or reduced levels of TGFβ transcriptional targets in *Sun2−/−* mice. WT and *Sun2−/−* Ct values were normalized to PPIA. Fold change of *Sun2−/−* mRNA levels was determined by calculating the 2^ΔΔCt^ after taking the mean of WT ΔCt values from n = 2 WT or *Sun2−/−* mice. Statistical significance determined by one-way ANOVA followed by Dunnett’s multiple comparisons test. P values are indicated in the graph. (E) qPCR analysis of left ventricular total RNA from 13 month old animals as described in (D).

### Disrupted TGFβ signaling and increased retention of MAN1 at the nuclear envelope in SUN2-null mice

One of the major signaling pathways known to drive fibrosis in the heart is the TGFβ/SMAD cascade (Massagué and Wotton, 2000; Massagué, 2012; Travers *et al.*, 2016). To determine the point at which the TGFβ signaling cascade (Massagué and Wotton, 2000; Massagué, 2012; Travers *et al.*, 2016) is disrupted in the absence of SUN2, we performed immunofluorescence on P4 WT and *Sun2−/−* cardiac sections using antibodies against phosphorylated SMAD2 (pSMAD2) (Figure 6A-B). Surprisingly, *Sun2−/−* cardiac tissue exhibited a striking increase in the nuclear localization of pSMAD2 in comparison to WT tissue (Figure 6A-B). These results raise the possibility that while the cytoplasmic aspects of TGFβ/SMAD signaling are increased in *Sun2−/−* cardiomyocytes, driving nuclear accumulation of pSMAD2, the nuclear aspects (transcription) are strongly abrogated in the absence of SUN2.

**Figure 6.**
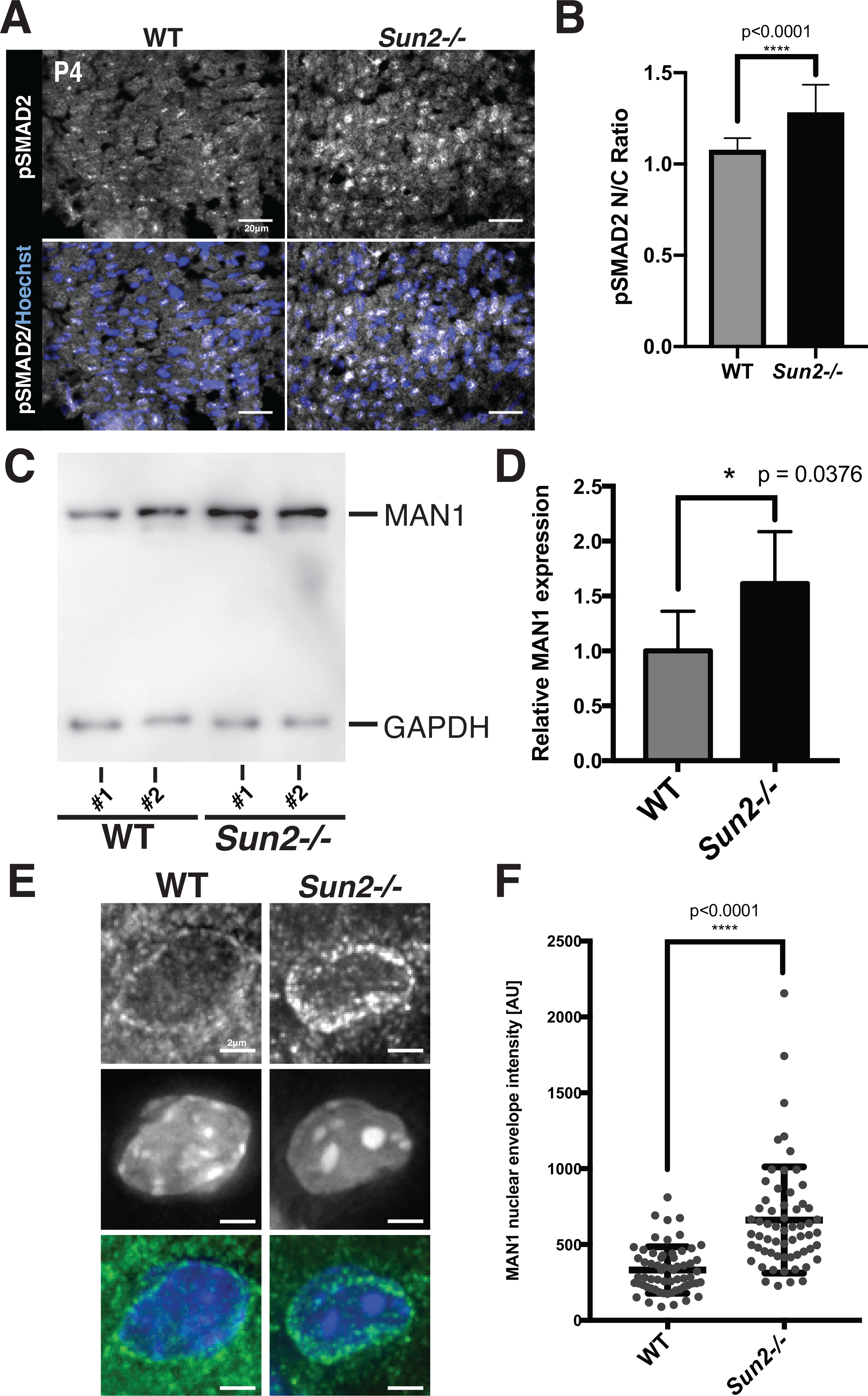
MAN1 levels at the nuclear envelope are increased in the *Sun2−/−* myocardium. (A) *Sun2−/−* tissue exhibits qualitatively increased pSMAD2 nuclear localization. Frozen P4 WT and *Sun2−/−* cardiac left ventricle tissue was sectioned and stained with antibodies against pSMAD2 (grey) and counterstained with Hoechst 33342 (blue). Scale bar = 20 om. (B) Enhanced pSMAD2 nuclear:cytoplasmic (N/C) ratio in *Sun2−/−* tissue. The relative maximum fluorescence intensity of the nuclear signal divided by the cytoplasmic signal for pSMAD2 in images obtained as in (A). n=503 cells for WT and n=545 cells for *Sun2−/−*. Statistical significance determined by unpaired *t* test with Welch’s correction. (C) Lysates from WT and *Sun2−/−* P50 mice were subjected to SDS-PAGE and immunoblotting with antibodies against MAN1 and GAPDH (as a loading control). (D) Quantification of densitometry analysis for MAN1 normalized to GAPDH. Data are represented as the mean ± SD for 4 WT or *Sun2−/−* mice. (E) Representative fluorescence micrographs of MAN1 staining in frozen sections of the P4 myocardium from WT or *Sun2−/−* mice. (F) Peak fluorescence intensity of MAN1 at the nuclear envelope determined by line-scan with local background subtraction. Plotted with the SD (n>50 nuclei). Statistical significance determined by unpaired *t* test with Welch’s correction.

As MAN1 is established to be both a lamin A/C-binding protein and a nuclear-localized, negative regulator of TGFβ, we next examined the levels and localization of MAN1 in the myocardium of *Sun2−/−* mice. AT P50, we found that total protein levels of MAN1 were moderately higher in *Sun2−/−* mice as detected by immunoblotting (Figure 6C-D). Further, in the myocardium of P50 SUN2-null mice we observed elevated levels of MAN1 by immunohistochemistry and quantitative analysis of fluorescence intensity across line profiles spanning the nuclear envelope (Figure 6E-F). Thus, while upstream events that drive pro-fibrotic pathway are likely stimulated in the absence of SUN2, mechanical signaling through the LINC complex may be required for nuclear pSMAD2 to act on its target genes, potentially by driving release of MAN1 from the nuclear lamina.

## Discussion

Here we demonstrate that a mouse model of SUN2 ablation develops cardiac hypertrophy, similar to existing mouse models of A-type lamin dysfunction. Surprisingly, we observe that the SUN2-null mouse, unlike laminopathic mouse models, is protected from the involvement of interstitial fibrosis, suggesting that although the pro-growth and pro-fibrotic pathways may be driven by the same upstream factors, these two outcomes can be genetically uncoupled, potentially through distinct impacts on the targeting of MAN1 to the nuclear lamina. These findings raise the exciting possibility that future studies will reveal how modulating the nuclear lamina can be leveraged to disrupt tissue fibrosis.

The earliest effects of SUN2 ablation on cardiac muscle manifest during development, consistent with the recent observation that the LINC complex nucleates developing sarcomeres at the nuclear envelope and promotes the maintenance of myofibril structure in *Drosophila* lateral transverse muscle (Auld and Folker, 2016), as well as the observation that overexpression of human Nesprin-1α2 in a zebrafish model leads to defects in cardiac development (Zhou *et al.*, 2017). Although we cannot rule out a direct role for LINC complexes in organized sarcomere assembly during the development of mammalian cardiac muscle, we favor a model in which altered regulation of the contractile-adhesion network underlies the observed sarcomere defects both early and late in life in *Sun2−/−* mice. Indeed, integrin-based adhesions at costameres play essential roles in patterning the sarcomere during cardiac myofibrillogenesis (Rhee *et al.*, 1994; Dabiri *et al.*, 1997; Hirschy *et al.*, 2006; Du *et al.*, 2008). Such a model is consistent with our previous demonstration that the LINC complex can influence cell junctions (Stewart *et al.*, 2015), and the apparent increase in the levels of β1-integrin in its active, ligand-bound conformation in *Sun2−/−* cardiac tissue (Figure 4A).

Further support for a potential foundational effect of increased contractility comes from our TEM analysis, which reveals that cardiomyocytes manifest with sarcomere defects reminiscent of those tied to hypercontractility (Lauritzen *et al.*, 2009) shortly after birth. Interestingly, a small molecular inhibitor of sarcomere power output in cardiomyocytes can be protective in an HCM-model, suggesting that increased cardiomyocyte contractility can drive cardiac hypertrophy (Green *et al.*, 2016). Thus, altered contractile properties of *Sun2−/−* cardiomyocytes could drive the LV hypertrophy we observe. How could SUN2 LINC complexes influence sarcomere contractile function? Beyond its direct interactions with the actin cytoskeleton, the LINC complex is implicated as a regulator of RhoA activity (Thakar *et al.*, 2017), interacts with other modulators of actin organization and function, such as the formin FHOD1 in fibroblasts (Kutscheidt *et al.*, 2014), and is known to influence actin dynamics (Lammerding *et al.*, 2004; Lee *et al.*, 2007; Hale *et al.*, 2008; Stewart-Hutchinson *et al.*, 2008; Khatau *et al.*, 2009; Luxton *et al.*, 2010b; Folker *et al.*, 2011; Kim *et al.*, 2012). However, it should be noted that while increased cardiac contractility at the organ-level is frequently observed in HCM, cell autonomous hypercontractility is not always observed in these disease models (Moore *et al.*, 2012). Recent work also suggests that the disease-associated D192G lamin A/C mutation may produce cytoskeletal and adhesion defects in neonatal rat cardiomyocytes (Lanzicher *et al.*, 2015), while a human cardiomyopathy patient with the point mutation G382V in *LMNA* exhibited defective plakoglobin localization to ICDs in the right ventricle (Quarta *et al.*, 2012). Whether cardiomyocytes in models of laminopathies exhibit altered adhesion function or contractile properties remains a critical future question.

Although it remains to be fully tested, we suggest that increased contractility in *Sun2−/−* mice could be responsible for the gain of β1-integrin engagement that we observe. Indeed, increased integrin activation can be driven by heightened intracellular actomyosin contractility, as has been shown for the integrin LFA-1 in migrating T cells (Nordenfelt *et al.*, 2016). Integrin engagement and subsequent activation of AKT signaling has also been shown to drive cardiac hypertrophy in mouse models (Reviewed in (Brancaccio *et al.*, 2006; Sequeira *et al.*, 2014)), and AKT activation is linked to elevated cardiomyocyte contractility (Kim *et al.*, 2003; Rota *et al.*, 2005; Cittadini *et al.*, 2006; Catalucci *et al.*, 2009), suggesting a potential connection between contractility, active β1-integrin levels and hypertrophy in *Sun2−/−* mice. Consistent with this model, we observe heightened levels of phosphorylated AKT at P50 in *Sun2−/−* mice (Figure 4B). Importantly, a similar transient AKT activation in adolescent mice followed by sustained AKT-mTOR downstream signaling has previously been described in the cardiac tissue of the lamin A/C H222P mutant mouse (Muchir *et al.*, 2007; Wu *et al.*, 2011; Choi *et al.*, 2012; Ramos *et al.*, 2012; Muchir *et al.*, 2012a; 2012b; Choi and Worman, 2013). The mechanisms by which loss of A-type lamin function leads to increased AKT and MAPK signaling, observed in numerous mouse models (Muchir *et al.*, 2007; Wu *et al.*, 2011; Choi *et al.*, 2012; Ramos *et al.*, 2012; Muchir *et al.*, 2012a; 2012b; Choi and Worman, 2013), has yet to be defined; our work suggests that further study of signaling from integrin-based adhesions may prove fruitful.

*LMNA−/−*, lamin A/C H222P and the *Sun2−/−* mouse models all share an increase in SMAD2 phosphorylation and nuclear localization (Chatzifrangkeskou *et al.*, 2016). In the context of lamin perturbation, this occurs coincident with the induction of TGFβ-associated gene expression and severe cardiac fibrosis (Chatzifrangkeskou *et al.*, 2016). By contrast, *Sun2−/−* mice display hypertrophy without fibrosis, mimicking physiological hypertrophy, which is characterized by hypertrophy with a lack of fibrosis or reactivation of fetal program genes (Maillet *et al.*, 2013). This is similar to what is observed during cardiac growth compensation in response to exercise and pregnancy (Frenzel *et al.*, 1988; Gonzalez *et al.*, 2007; Chung *et al.*, 2012), or in models of AMP-insensitive “activated” AMPK mutations (Hinson *et al.*, 2016). One model is that SUN2 is a required component of a pathway that “licenses” nuclear pSMAD2 to act on its target genes. Our observation that MAN1 accumulates at the nuclear lamina in the absence of SUN2 suggests that its repression of TGFβ/SMAD signaling is alleviated in contexts where the LINC complex is under tension.

Importantly, while treatment of mouse models of lamin dysfunction with the TOR inhibitor rapamycin positively influences heart function and ameliorates cardiac hypertrophy (Muchir *et al.*, 2007; Wu *et al.*, 2011; Choi *et al.*, 2012; Ramos *et al.*, 2012; Muchir *et al.*, 2012a; 2012b; Choi and Worman, 2013), it does not reverse the coincident fibrosis, highlighting that these pathways can be uncoupled. As human laminopathy patients typically display left ventricular dilation coupled with fibrosis (Ostlund *et al.*, 2001; Meinke *et al.*, 2014; Stroud *et al.*, 2014), identifying the mechanisms that drive pro-fibrotic signaling remains a critical need. Further study of the nuclear aspect of this signaling cascade through modulating MAN1 may identify new approaches for intervening in pathological fibrosis.

## Acknowledgements

This work was supported by American Heart Association Predoctoral Fellowship #16PRE27460000 (RMS), an Outstanding Early Investigator Award from the Ludwig Family Foundation (MCK) and R01 GM129308-01 from the NIH/NIGMS (MCK). We thank Dr. Lawrence Young and Dr. Stuart Campbell of Yale University for valuable discussions, Morven Graham, Nicole Mikush, and Kevin Su for their technical expertise, and Dr. Brian Burke for the kind gift of the antibody to MAN1 for staining of tissue. The authors declare no competing financial interests.

## Author Contributions

R.M.S. designed and carried out experiments, analyzed data, prepared the figures and wrote the manuscript. E.C.R. maintained the mouse colony and carried out experiments, M.C.K. designed the experiments, analyzed the data, and edited the figures and manuscript. All authors approved of the final manuscript.

## Materials and Methods

### Mouse breading and care

All animal care and experimental procedures were conducted in accord with requirements approved by the Institutional Animal Care and Use Committee of Yale University. *Sun2^−/−^* (strain B6;129S6-*Sun2^tm1Mhan^*/J) and C57BL/6 WT mice were obtained from Jackson ImmunoResearch Laboratories, Inc. *Sun2^−/−^* mice were previously generated through the replacement of exons 11–16 and part of exon 17 with a neomycin resistance cassette (Lei *et al.*, 2009).

### Mouse tissue isolation, histology, and immunofluorescence staining

#### Cardiac isolations

WT and *Sun2−/−* murine hearts were isolated from P4 or P45/P47/P50 mice, perfused with PBS, and left ventricular tissue was either frozen in O.C.T. compound (Sakura, Tissue-Tek) at −80°C for cryo-sectioning or fixed in 10% neutral buffered formalin and embedded in paraffin in collaboration with the Yale University Developmental Histology Facility in the Department of Pathology. Frozen samples were sectioned using a cryostat (CM3050S; Leica). For histological analyses, 10-μm sections were cut and stained with hematoxylin and eosin. For immunofluorescence, 6- or 8-μm sections were fixed in 4% formaldehyde or 4% paraformaldehyde at RT for 10 minutes.

#### Immunofluorescence staining

For immunostaining with nonmouse primary antibodies, tissue sections were blocked in gelatin block (2.5% normal goat serum, 1% BSA, 2% gelatin, and 0.25% Triton X-100 in PBS) at RT for 1 h and incubated with the following primary antibodies overnight at 4°C: SUN2 (1:100; rabbit; Abcam ab124916), total β1-integrin (1:50; rat; Novus Biologicals NBP1-4323, clone KMI6), conformational epitope β1-integrin (1:50; rat; BD Pharmingen #550531, clone 9EG7), paxillin (1:100; rabbit; Abcam ab32084, clone Y113), or phosphorylated SMAD2 (1:100, rabbit, Cell Signaling #8828, clone D27F4). For immunostaining with the mouse primary antibodies desmoplakin I/II (1:50; mouse; Abcam ab16434, clone 2Q400), β-Catenin (1:100; mouse; BD Transduction Labs #610153), and anti-MAN1 antibody (kind gift from Brian Burke), the Mouse-on-Mouse (MOM) Immunodetection kit Blocking Reagent and Protein Diluent (Vector Laboratories) were used according to manufacturer’s instructions. Sections were subsequently washed in multiple changes of PBS and incubated with fluorescent dye–conjugated secondary antibodies (1:1,000; mouse, rat, or rabbit; Alexa Fluor; Life Technologies) in gel block or MOM-containing gel block when appropriate. Costaining with Hoechst 33342 (Thermo Fisher Scientific; 1:2,000), Alexa Fluor 594-conjugated wheat germ agglutinin (Life Technologies; 1:2,000), and/or Alexa Fluor 594-conjugated phalloidin (Invitrogen #A12381; 1:40) was performed when indicated. Sections were mounted using Fluoromount-G mounting medium (SouthernBiotech).

### Transmission electron microscopy

TEM was performed in the Yale School of Medicine Center for Cellular and Molecular Imaging Electron Microscopy core facility. Left ventricular cardiac tissue from P4 and P45/P47 WT and *Sun2^−/−^* mice were isolated and processed for TEM; three mice were examined for each genotype at each time point. Tissue blocks were fixed in 2.5% glutaraldehyde/2% paraformaldehyde in 0.1 M sodium cacodylate buffer, pH 7.4, for 30 min at RT and 1.5 h at 4°C. The samples were rinsed in sodium cacodylate buffer and were postfixed in 1% osmium tetroxide for 1 h. The samples were rinsed and en bloc stained in aqueous 2% uranyl acetate for 1 h followed by rinsing, dehydrating in an ethanol series to 100%, rinsing in 100% propylene oxide, infiltrating with EMbed 812 (Electron Microscopy Sciences) resin, and baking overnight at 60°C. Hardened blocks were cut using an ultramicrotome (UltraCut UC7; Leica). Ultrathin 60-nm sections were collected and stained using 2% uranyl acetate and lead citrate for transmission microscopy. Carbon-coated grids were viewed on a transmission electron microscope (Tecnai BioTWIN; FEI) at 80 kV. Images were taken using a CCD camera (Morada; Olympus) and iTEM (Olympus) software.

### Imaging and image analysis

#### Immunofluorescence imaging

Whole slides of Hematoxylin and eosin-stained or trichrome-stained cardiac sections were scanned using an Aperio Digital Pathology Scanner at 40x magnification in collaboration with the Yale University Pathology Digital Imaging in the Department of Pathology. Cardiac sections processed for immunofluorescence were imaged on a Leica SP5 confocal microscope or a widefield deconvolution microscope (DeltaVision; Applied Precision/GE Healthcare) with a charge-coupled device (CCD) camera (CoolSNAP K4; Photometrics) and SoftWoRx software. All images acquired on the DeltaVision microscope were deconvolved using the Deconvolve tool (constrained iterative deconvolution) in SoftWoRx software. The DeltaVision microscope was equipped with an oil Plan Apochromat N 60×/1.42 NA objective (Olympus). In all cases, images were analyzed using Fiji software (ImageJ 1.48d; National Institutes of Health) as indicated.

#### Analysis of cardiomyocyte size

Transverse midventricular sections of WT and *Sun2−/−* murine hearts were cut and stained with antibodies against laminin in collaboration with the Yale University Developmental Histology Facility in the Department of Pathology. The cross-sectional area of between 86 – 198 left ventricular papillary muscle cells was measured by manually outlining individual cells and using the measure tool in Fiji (ImageJ 1.51j; National Institutes of Health)(Schindelin *et al.*, 2012). All cells in a given field were measured and only fields with circular blood vessels – indicating that the adjacent cells were sectioned orthogonal to the plane – were included.

#### Analysis of fibrosis

Transverse midventricular sections of WT and *Sun2−/−* murine hearts were cut and stained with Masson’s trichrome stain in collaboration with the Yale University Developmental Histology Facility in the Department of Pathology. Whole slides were scanned using an Aperio Digital Pathology Scanner at 40x magnification in collaboration with the Yale University Pathology Digital Imaging in the Department of Pathology. The ImageScope 12.2 Positive Pixel Count algorithm (Aperio Technologies, Inc.) was used to identify blue-colored collagen fibers in the tissue samples. Color hue and saturation values (hue value of 0.62, hue width of 0.40, and color saturation threshold of 0.005) were optimized before the analysis. Five or more fields of interstitial tissue were assessed for each of 3 mice per genotype.

#### Analysis of sarcomere length

TEM micrographs of P50 WT and *Sun2−/−* ventricular tissue were acquired as described. The sarcomere length, defined as the distance from the center of one Z-band to the center of a successive Z-band, was measured in Fiji for between 94 – 235 sarcomeres for 3 mice of each genotype. All sarcomeres with visible Z-bands in a given field were measured.

#### Display of β1-integrin fluorescence intensity

Images of WT and *Sun2−/−* ventricular tissue stained with antibodies against the ligand-bound β1-integrin conformation were acquired on the Leica SP5 microscope using the same acquisition settings between WT and *Sun2−/−* samples. WT and *Sun2−/−* images were scaled to one another and pseudo-colored in Fiji using the Fire LUT, converting differences in pixel intensity to differences in display color; on this scale, lighter colors correspond to greater fluorescence intensity while darker colors correspond to lower fluorescence intensity.

#### pSMAD2 nuclear:cytoplasmic intensity ratio measurements

Using images as in Figure 6A, nuclei were segmented by thresholding the Hoechst signal, binarizing, and creating individual nuclear masks using the Analyze Particles function in ImageJ. The resulting masks were used to measure the nuclear pSMAD2 fluorescence intensity maxima following background subtraction. The masks were converted to a band of 1.5 pixels to measure perinuclear cytoplasmic intensity maxima. The nuclear:cytoplasmic intensity ratio was calculated as the nuclear intensity maximum divided by cytoplasmic intensity maximum.

#### MAN1 fluorescence intensity measurements

Using images as in Figure 6E, line profiles bisecting the nucleus of equal length were taken (>50 per sample) in ImageJ. The local intensity maxima at the nuclear envelope after subtracting the mean intensity across the entire line profile was then calculated.

### Western blotting

Following heart isolation and PBS perfusion, ventricular cardiac tissue was isolated from 13 month-old WT and *Sun2−/−* mice or P45/P47 WT and *Sun2−/−* mice, minced, and homogenized using a Polytron PT 1200 E homogenizer (Kinematica AG) in radioimmunoprecipitation lysis buffer (50 mM Tris-Cl, pH 7.4, 1 mM EDTA, 1% NP-40 alternative, 0.1% Na deoxycholate, 0.1% SDS, 150 mM NaCl, protease inhibitor cocktail (Sigma-Aldrich). The insoluble debris was removed by centrifugation for 30 min at 14,000 rpm in a 4°C tabletop microcentrifuge (Sorvall Legend Micro 21R; Thermo Fisher Scientific). Extracted protein samples were diluted in SDS-PAGE sample buffer and separated using a 7.5% polyacrylamide gel. The separated proteins were transferred to a nitrocellulose membrane using a transfer system (Bio-Rad Laboratories) and subjected to Ponceau S (Sigma) staining to observe total protein loading. Membranes were subsequently blocked in 10% nonfat milk (Omniblok; American Bioanalytical) or, if blotting with phospho-antibodies, 5% BSA (AmericanBio) in TBST (TBS with Tween 20) for 1 h at RT and incubated with AKT (1:1000, rabbit, Cell Signaling 9272), pAKT (1:1000, rabbit, Cell Signaling 4060 clone D9E), S6 (1:5000, rabbit, Cell Signaling 2217 clone5G10), pS6 (1:5000, rabbit, Cell Signaling 2211 Ser235/236), ERK2 (1:200, rabbit, Santa Cruz SC-154), pERK1/2 (1:300, mouse, Cell Signaling 9101 Thr202/204), and MAN1/LEMD3 (1:1000, rabbit, ProSci 6603) primary antibodies at 4°C overnight. The membranes were washed for 15 min in TBST and then incubated for 1 h with HRP-conjugated secondary antibodies (1:5,000; Pierce Antibodies; Thermo Fisher Scientific) in TBST at RT. After washes with TBST, the membranes were incubated with HRP chemiluminescent substrate (SuperSignal West Femto Chemiluminescent Substrate; Thermo Fisher Scientific) for 5 min and then imaged using a VersaDoc imaging system (Bio-Rad Laboratories).

### qPCR

Following heart isolation and PBS perfusion, ventricular cardiac tissue was isolated from 13-month-old WT and *Sun2−/−* mice or P54/P55 WT and *Sun2−/−* mice, minced, and homogenized in TRIzol (Invitrogen). Total RNA was isolated using the RNeasy kit (QIAGEN) according to manufacturer’s instructions. To remove genomic DNA, total RNA was digested with DNase I (Roche) according to manufacturer’s instructions and repurified using the RNeasy kit (QIAGEN). To generate cDNA, equal amounts of digested total RNA (1 μg) were added to SuperScript III Reverse Transcriptase (Invitrogen) reactions using Random Primer 9 (New England Biolabs).

Quantitative real-time PCR was conducted with a Bio-Rad CFX96 Real-Time System using the iQ SYBR Green Supermix (Bio-Rad) for 39 cycles. Primers used in these experiments were: ANP forward: CAAGATGCAGAAGCTGCTGG and reverse: GTGCTGCCTTGAGACCGAAGG; BNP forward: GTTTGGGCTGTAACGCACTGA and reverse: GAAAGAGACCCAGGCAGAGTCA; Skeletal α-actin forward: CATGAAGATCAAGATCATCGC and reverse: CTGGAAGGTGGACAGCGAGGC; Serca2 forward: GTGTGGCAGGAAAGAAATGC and reverse: CCAGGAACTATGTCTTTAGC; FN1 forward: TTCAAGTGTGATCCCCATGAAG and reverse: CAGGTCTACGGCAGTTGTCA; COL1A1 forward: CTGGCGGTTCAGGTCCAAT and reverse: TTCAGGCAATCCAGAGC; COL3A1 forward: CTGTAACATGGAAACTGGGGAAA and ereverse: CCATAGCTGAACTGAAAACCACC; POSTN forward: CCTGCCCTTATATGCTCTGCT and reverse: AAACATGGTCAATAGGCATCACT; LTBP2 forward: GCTCACCGGGAGAAATGTCTG and reverse: CAGGTTTGATACAGTGGTTGGT; LOXL1 forward: GAGTGCTATTGCGCTTCCC and reverse: GGTTGCCGAAGTCACAGGT; FLNA forward: GGCTACGGTGGGCTTAGTC and reverse: GTGGGACAGTAGGTGACCCT; DES forward: TACACCTGCGAGATTGATGC and reverse: ACATCCAAGGCCATCTTCAC; GAPDH forward: CGTAGACAAAATGGTGAAGGTCGG and reverse: AAGCAGTTGGTGGTGCAGGATG; and PPIA forward: GAGCTGTTTGCAGACAAAGTTC and reverse: CCCTGGCACATGAATCCTGG. PCR product levels were normalized to peptidylprolyl isomerase A (PPIA) or glyceraldehyde 3-phosphate dehydrogenase (GAPDH) mRNA levels.

## Abbreviation

LINC complex: linker of Nucleoskeleton and Cytoskeleton
ICD: intercalated disc
LV: left ventricle
HCM: hypertrophic cardiomyopathy
qPCR: quantitative polymerase chain reaction
TEM: transmission electron microscopy

## References

Arbustini, E. et al. (2002). Autosomal dominant dilated cardiomyopathy with atrioventricular block: a lamin A/C defect-related disease. J. Am. Coll. Cardiol. 39, 981–990.

Auld, A. L., and Folker, E. S. (2016). Nucleus-dependent sarcomere assembly is mediated by the LINC complex. Mol. Biol. Cell 27, 2351–2359.

Balasubramanian, S., and Kuppuswamy, D. (2003). RGD-containing peptides activate S6K1 through beta3 integrin in adult cardiac muscle cells. J. Biol. Chem. 278, 42214–42224.

Bazzoni, G., Shih, D. T., Buck, C. A., and Hemler, M. E. (1995). Monoclonal antibody 9EG7 defines a novel beta 1 integrin epitope induced by soluble ligand and manganese, but inhibited by calcium. J. Biol. Chem. 270, 25570–25577.

Brancaccio, M., Hirsch, E., Notte, A., Selvetella, G., Lembo, G., and Tarone, G. (2006). Integrin signalling: the tug-of-war in heart hypertrophy. Cardiovasc. Res. 70, 422–433.

Cahill, T. J., Ashrafian, H., and Watkins, H. (2013). Genetic cardiomyopathies causing heart failure. Circ. Res. 113, 660–675.

Catalucci, D., Latronico, M. V. G., Ceci, M., Rusconi, F., Young, H. S., Gallo, P., Santonastasi, M., Bellacosa, A., Brown, J. H., and Condorelli, G. (2009). Akt increases sarcoplasmic reticulum Ca2+ cycling by direct phosphorylation of phospholamban at Thr17. J. Biol. Chem. 284, 28180–28187.

Chambers, D. M., Moretti, L., Zhang, J. J., Cooper, S. W., Chambers, D. M., Santangelo, P. J., and Barker, T. H. (2018). LEM domain-containing protein 3 antagonizes TGFβ-SMAD2/3 signaling in a stiffness-dependent manner in both the nucleus and cytosol. J. Biol. Chem. 293, 15867–15886.

Chang, W., Worman, H. J., and Gundersen, G. G. (2015). Accessorizing and anchoring the LINC complex for multifunctionality. J. Cell Biol. 208, 11–22.

Chatzifrangkeskou, M. et al. (2016). ERK1/2 directly acts on CTGF/CCN2 expression to mediate myocardial fibrosis in cardiomyopathy caused by mutations in the lamin A/C gene. Hum. Mol. Genet. 25, 2220–2233.

Choi, J. C., and Worman, H. J. (2013). Reactivation of autophagy ameliorates LMNA cardiomyopathy. Autophagy 9, 110–111.

Choi, J. C., Muchir, A., Wu, W., Iwata, S., Homma, S., Morrow, J. P., and Worman, H. J. (2012). Temsirolimus activates autophagy and ameliorates cardiomyopathy caused by lamin A/C gene mutation. Sci Transl Med 4, 144ra102–144ra102.

Chung, E., Yeung, F., and Leinwand, L. A. (2012). Akt and MAPK signaling mediate pregnancy-induced cardiac adaptation. J. Appl. Physiol. 112, 1564–1575.

Cittadini, A. et al. (2006). Adenoviral gene transfer of Akt enhances myocardial contractility and intracellular calcium handling. Gene Ther. 13, 8–19.

Dabiri, G. A., Turnacioglu, K. K., Sanger, J. M., and Sanger, J. W. (1997). Myofibrillogenesis visualized in living embryonic cardiomyocytes. Proc Natl Acad Sci U S A 94, 9493–9498.

Davidson, P. M., and Lammerding, J. (2014). Broken nuclei--lamins, nuclear mechanics, and disease. Trends Cell Biol. 24, 247–256.

Delmar, M., and McKenna, W. J. (2010). The cardiac desmosome and arrhythmogenic cardiomyopathies: from gene to disease. Circ. Res. 107, 700–714.

Dowling, J. J., Gibbs, E., Russell, M., Goldman, D., Minarcik, J., Golden, J. A., and Feldman, E. L. (2008). Kindlin-2 is an essential component of intercalated discs and is required for vertebrate cardiac structure and function. Circ. Res. 102, 423–431.

Du, A., Sanger, J. M., and Sanger, J. W. (2008). Cardiac myofibrillogenesis inside intact embryonic hearts. Dev. Biol. 318, 236–246.

Folker, E. S., Ostlund, C., Luxton, G. W. G., Worman, H. J., and Gundersen, G. G. (2011). Lamin A variants that cause striated muscle disease are defective in anchoring transmembrane actin-associated nuclear lines for nuclear movement. Proc Natl Acad Sci U S A 108, 131–136.

Franchini, K. G., Torsoni, A. S., Soares, P. H., and Saad, M. J. (2000). Early activation of the multicomponent signaling complex associated with focal adhesion kinase induced by pressure overload in the rat heart. Circ. Res. 87, 558–565.

Frenzel, H., Schwartzkopff, B., Höltermann, W., Schnürch, H. G., Novi, A., and Hort, W. (1988). Regression of cardiac hypertrophy: morphometric and biochemical studies in rat heart after swimming training. J. Mol. Cell. Cardiol. 20, 737–751.

Frock, R. L. et al. (2012). Cardiomyocyte-specific expression of lamin a improves cardiac function in Lmna−/− mice. PLoS One 7, e42918.

Gomez, E. W., Chen, Q. K., Gjorevski, N., and Nelson, C. M. (2010). Tissue geometry patterns epithelial-mesenchymal transition via intercellular mechanotransduction. Journal of Cellular Biochemistry 110, 44–51.

Gonzalez, A. M. D., Osorio, J. C., Manlhiot, C., Gruber, D., Homma, S., and Mital, S. (2007). Hypertrophy signaling during peripartum cardiac remodeling. Am. J. Physiol. Heart Circ. Physiol. 293, H3008–H3013.

Green, E. M. et al. (2016). A small-molecule inhibitor of sarcomere contractility suppresses hypertrophic cardiomyopathy in mice. Science 351, 617–621.

Gupta, P. et al. (2010). Genetic and ultrastructural studies in dilated cardiomyopathy patients: a large deletion in the lamin A/C gene is associated with cardiomyocyte nuclear envelope disruption. Basic Res. Cardiol. 105, 365–377.

Hale, C. M., Shrestha, A. L., Khatau, S. B., Stewart-Hutchinson, P. J., Hernandez, L., Stewart, C. L., Hodzic, D., and Wirtz, D. (2008). Dysfunctional connections between the nucleus and the actin and microtubule networks in laminopathic models. Biophys. J. 95, 5462–5475.

Hatch, E. M., and Hetzer, M. W. (2016). Nuclear envelope rupture is induced by actin-based nucleus confinement. J. Cell Biol. 215, 27–36.

Hinson, J. T. et al. (2016). Integrative Analysis of PRKAG2 Cardiomyopathy iPS and Microtissue Models Identifies AMPK as a Regulator of Metabolism, Survival, and Fibrosis. Cell Rep 17, 3292–3304.

Hinz, B. (2015). The extracellular matrix and transforming growth factor-β1: Tale of a strained relationship. Matrix Biol. 47, 54–65.

Hirschy, A., Schatzmann, F., Ehler, E., and Perriard, J.-C. (2006). Establishment of cardiac cytoarchitecture in the developing mouse heart. Dev. Biol. 289, 430–441.

Ho, C. Y. et al. (2010). Myocardial fibrosis as an early manifestation of hypertrophic cardiomyopathy. N. Engl. J. Med. 363, 552–563.

Iijima, Y., Laser, M., Shiraishi, H., Willey, C. D., Sundaravadivel, B., Xu, L., McDermott, P. J., and Kuppuswamy, D. (2002). c-Raf/MEK/ERK pathway controls protein kinase C-mediated p70S6K activation in adult cardiac muscle cells. J. Biol. Chem. 277, 23065–23075.

Ishimura, A., Ng, J. K., Taira, M., Young, S. G., and Osada, S.-I. (2006). Man1, an inner nuclear membrane protein, regulates vascular remodeling by modulating transforming growth factor beta signaling. Development 133, 3919–3928.

Israeli-Rosenberg, S., Manso, A. M., Okada, H., and Ross, R. S. (2014). Integrins and integrin-associated proteins in the cardiac myocyte. Circ. Res. 114, 572–586.

Khatau, S. B. et al. (2012). The distinct roles of the nucleus and nucleus-cytoskeleton connections in three-dimensional cell migration. Sci Rep 2, 488.

Khatau, S. B., Hale, C. M., Stewart-Hutchinson, P. J., Patel, M. S., Stewart, C. L., Searson, P. C., Hodzic, D., and Wirtz, D. (2009). A perinuclear actin cap regulates nuclear shape. Proc Natl Acad Sci U S A 106, 19017–19022.

Kim, D.-H., Khatau, S. B., Feng, Y., Walcott, S., Sun, S. X., Longmore, G. D., and Wirtz, D. (2012). Actin cap associated focal adhesions and their distinct role in cellular mechanosensing. Sci Rep 2, 555.

Kim, J. B., Porreca, G. J., Song, L., Greenway, S. C., Gorham, J. M., Church, G. M., Seidman, C. E., and Seidman, J. G. (2007). Polony multiplex analysis of gene expression (PMAGE) in mouse hypertrophic cardiomyopathy. Science 316, 1481–1484.

Kim, Y.-K. et al. (2003). Mechanism of enhanced cardiac function in mice with hypertrophy induced by overexpressed Akt. J. Biol. Chem. 278, 47622–47628.

Konieczny, P., Fuchs, P., Reipert, S., Kunz, W. S., Zeöld, A., Fischer, I., Paulin, D., Schröder, R., and Wiche, G. (2008). Myofiber integrity depends on desmin network targeting to Z-disks and costameres via distinct plectin isoforms. J. Cell Biol. 181, 667–681.

Kutscheidt, S., Zhu, R., Antoku, S., Luxton, G. W. G., Stagljar, I., Fackler, O. T., and Gundersen, G. G. (2014). FHOD1 interaction with nesprin-2G mediates TAN line formation and nuclear movement. Nat. Cell Biol. 16, 708–715.

Lammerding, J., Schulze, P. C., Takahashi, T., Kozlov, S., Sullivan, T., Kamm, R. D., Stewart, C. L., and Lee, R. T. (2004). Lamin A/C deficiency causes defective nuclear mechanics and mechanotransduction. J. Clin. Invest. 113, 370–378.

Lanzicher, T., Martinelli, V., Puzzi, L., Del Favero, G., Codan, B., Long, C. S., Mestroni, L., Taylor, M. R. G., and Sbaizero, O. (2015). The Cardiomyopathy Lamin A/C D192G Mutation Disrupts Whole-Cell Biomechanics in Cardiomyocytes as Measured by Atomic Force Microscopy Loading-Unloading Curve Analysis. Sci Rep 5, 13388.

Laser, M., Willey, C. D., Jiang, W., Cooper, G., Menick, D. R., Zile, M. R., and Kuppuswamy, D. (2000). Integrin activation and focal complex formation in cardiac hypertrophy. J. Biol. Chem. 275, 35624–35630.

Lauritzen, F., Paulsen, G., Raastad, T., Bergersen, L. H., and Owe, S. G. (2009). Gross ultrastructural changes and necrotic fiber segments in elbow flexor muscles after maximal voluntary eccentric action in humans. J. Appl. Physiol. 107, 1923–1934.

Le Dour, C., Wu, W., Béréziat, V., Capeau, J., Vigouroux, C., and Worman, H. J. (2017). Extracellular matrix remodeling and transforming growth factor-β signaling abnormalities induced by lamin A/C variants that cause lipodystrophy. J. Lipid Res. 58, 151–163.

Lee, J. S. H., Hale, C. M., Panorchan, P., Khatau, S. B., George, J. P., Tseng, Y., Stewart, C. L., Hodzic, D., and Wirtz, D. (2007). Nuclear lamin A/C deficiency induces defects in cell mechanics, polarization, and migration. Biophys. J. 93, 2542–2552.

Lei, K., Zhang, X., Ding, X., Guo, X., Chen, M., Zhu, B., Xu, T., Zhuang, Y., Xu, R., and Han, M. (2009). SUN1 and SUN2 play critical but partially redundant roles in anchoring nuclei in skeletal muscle cells in mice. Proc Natl Acad Sci U S A 106, 10207–10212.

Lenter, M., Uhlig, H., Hamann, A., Jenö, P., Imhof, B., and Vestweber, D. (1993). A monoclonal antibody against an activation epitope on mouse integrin chain beta 1 blocks adhesion of lymphocytes to the endothelial integrin alpha 6 beta 1. Proc Natl Acad Sci U S A 90, 9051–9055.

Liao, C.-Y. et al. (2016). Rapamycin Reverses Metabolic Deficits in Lamin A/C-Deficient Mice. Cell Rep 17, 2542–2552.

Lin, F., Morrison, J. M., Wu, W., and Worman, H. J. (2005). MAN1, an integral protein of the inner nuclear membrane, binds Smad2 and Smad3 and antagonizes transforming growth factor-beta signaling. Hum. Mol. Genet. 14, 437–445.

Long, J. B., Bagonis, M., Lowery, L. A., Lee, H., Danuser, G., and Van Vactor, D. (2013). Multiparametric analysis of CLASP-interacting protein functions during interphase microtubule dynamics. Mol. Cell. Biol. 33, 1528–1545.

Lowey, S. (2002). Functional consequences of mutations in the myosin heavy chain at sites implicated in familial hypertrophic cardiomyopathy. Trends Cardiovasc. Med. 12, 348–354.

Luxton, G. W. G., Gomes, E. R., Folker, E. S., Vintinner, E., and Gundersen, G. G. (2010a). Linear arrays of nuclear envelope proteins harness retrograde actin flow for nuclear movement. Science 329, 956–959.

Luxton, G. W. G., Gomes, E. R., Folker, E. S., Vintinner, E., and Gundersen, G. G. (2010b). Linear Arrays of Nuclear Envelope Proteins Harness Retrograde Actin Flow for Nuclear Movement. Science 329, 956–959.

MacKenna, D. A., Dolfi, F., Vuori, K., and Ruoslahti, E. (1998). Extracellular signal-regulated kinase and c-Jun NH2-terminal kinase activation by mechanical stretch is integrin-dependent and matrix-specific in rat cardiac fibroblasts. J. Clin. Invest. 101, 301–310.

Maillet, M., van Berlo, J. H., and Molkentin, J. D. (2013). Molecular basis of physiological heart growth: fundamental concepts and new players. Nat. Rev. Mol. Cell Biol. 14, 38–48.

Massagué, J. (2012). TGFβ signalling in context. Nat. Rev. Mol. Cell Biol. 13, 616–630.

Massagué, J., and Wotton, D. (2000). Transcriptional control by the TGF-beta/Smad signaling system. Embo J. 19, 1745–1754.

McCain, M. L., and Parker, K. K. (2011). Mechanotransduction: the role of mechanical stress, myocyte shape, and cytoskeletal architecture on cardiac function. Pflugers Arch. 462, 89–104.

McCain, M. L., Lee, H., Aratyn-Schaus, Y., Kléber, A. G., and Parker, K. K. (2012). Cooperative coupling of cell-matrix and cell-cell adhesions in cardiac muscle. Proc Natl Acad Sci U S A 109, 9881–9886.

Meinke, P. et al. (2014). Muscular dystrophy-associated SUN1 and SUN2 variants disrupt nuclear-cytoskeletal connections and myonuclear organization. PLoS Genet. 10, e1004605.

Michele, D. E., Albayya, F. P., and Metzger, J. M. (1999). Direct, convergent hypersensitivity of calcium-activated force generation produced by hypertrophic cardiomyopathy mutant alpha-tropomyosins in adult cardiac myocytes. Nat. Med. 5, 1413–1417.

Moore, J. R., Leinwand, L., and Warshaw, D. M. (2012). Understanding cardiomyopathy phenotypes based on the functional impact of mutations in the myosin motor. Circ. Res. 111, 375–385.

Mounkes, L. C., Kozlov, S. V., Rottman, J. N., and Stewart, C. L. (2005). Expression of an LMNA-N195K variant of A-type lamins results in cardiac conduction defects and death in mice. Hum. Mol. Genet. 14, 2167–2180.

Muchir, A., Pavlidis, P., Decostre, V., Herron, A. J., Arimura, T., Bonne, G., and Worman, H. J. (2007). Activation of MAPK pathways links LMNA mutations to cardiomyopathy in Emery-Dreifuss muscular dystrophy. J. Clin. Invest. 117, 1282–1293.

Muchir, A., Reilly, S. A., Wu, W., Iwata, S., Homma, S., Bonne, G., and Worman, H. J. (2012a). Treatment with selumetinib preserves cardiac function and improves survival in cardiomyopathy caused by mutation in the lamin A/C gene. Cardiovasc. Res. 93, 311–319.

Muchir, A., Wu, W., Choi, J. C., Iwata, S., Morrow, J., Homma, S., and Worman, H. J. (2012b). Abnormal p38α mitogen-activated protein kinase signaling in dilated cardiomyopathy caused by lamin A/C gene mutation. Hum. Mol. Genet. 21, 4325–4333.

Myat, M. M., Rashmi, R. N., Manna, D., Xu, N., Patel, U., Galiano, M., Zielinski, K., Lam, A., and Welte, M. A. (2015). Drosophila KASH-domain protein Klarsicht regulates microtubule stability and integrin receptor localization during collective cell migration. Dev. Biol. 407, 103–114.

Nordenfelt, P., Elliott, H. L., and Springer, T. A. (2016). Coordinated integrin activation by actin-dependent force during T-cell migration. Nat Commun 7, 13119.

O’Connor, J. W., Riley, P. N., Nalluri, S. M., Ashar, P. K., and Gomez, E. W. (2015). Matrix Rigidity Mediates TGFβ1-Induced Epithelial-Myofibroblast Transition by Controlling Cytoskeletal Organization and MRTF-A Localization. J. Cell. Physiol. 230, 1829–1839.

Olive, M. et al. (2010). Cardiovascular pathology in Hutchinson-Gilford progeria: correlation with the vascular pathology of aging. Arterioscler. Thromb. Vasc. Biol. 30, 2301–2309.

Ostlund, C., Bonne, G., Schwartz, K., and Worman, H. J. (2001). Properties of lamin A mutants found in Emery-Dreifuss muscular dystrophy, cardiomyopathy and Dunnigan-type partial lipodystrophy. J. Cell. Sci. 114, 4435–4445.

Ostlund, C., Sullivan, T., Stewart, C. L., and Worman, H. J. (2006). Dependence of diffusional mobility of integral inner nuclear membrane proteins on A-type lamins. Biochemistry 45, 1374–1382.

Quarta, G., Syrris, P., Ashworth, M., Jenkins, S., Zuborne Alapi, K., Morgan, J., Muir, A., Pantazis, A., McKenna, W. J., and Elliott, P. M. (2012). Mutations in the Lamin A/C gene mimic arrhythmogenic right ventricular cardiomyopathy. Eur. Heart J. 33, 1128–1136.

Raju, G. P., Dimova, N., Klein, P. S., and Huang, H.-C. (2003). SANE, a novel LEM domain protein, regulates bone morphogenetic protein signaling through interaction with Smad1. J. Biol. Chem. 278, 428–437.

Ramos, F. J. et al. (2012). Rapamycin reverses elevated mTORC1 signaling in lamin A/C-deficient mice, rescues cardiac and skeletal muscle function, and extends survival. Sci Transl Med 4, 144ra103–144ra103.

Rhee, D., Sanger, J. M., and Sanger, J. W. (1994). The premyofibril: evidence for its role in myofibrillogenesis. Cell Motility and the Cytoskeleton 28, 1–24.

Rota, M. et al. (2005). Nuclear targeting of Akt enhances ventricular function and myocyte contractility. Circ. Res. 97, 1332–1341.

Schindelin, J. et al. (2012). Fiji: an open-source platform for biological-image analysis. Nat. Methods 9, 676–682.

Sequeira, V., Nijenkamp, L. L. A. M., Regan, J. A., and van der Velden, J. (2014). The physiological role of cardiac cytoskeleton and its alterations in heart failure. Biochim. Biophys. Acta 1838, 700–722.

Sheikh, F., Ross, R. S., and Chen, J. (2009). Cell-cell connection to cardiac disease. Trends Cardiovasc. Med. 19, 182–190.

Shiojima, I., and Walsh, K. (2006). Regulation of cardiac growth and coronary angiogenesis by the Akt/PKB signaling pathway. Genes Dev. 20, 3347–3365.

Sosa, B. A., Rothballer, A., Kutay, U., and Schwartz, T. U. (2012). LINC complexes form by binding of three KASH peptides to domain interfaces of trimeric SUN proteins. Cell 149, 1035–1047.

Stewart, R. M., Zubek, A. E., Rosowski, K. A., Schreiner, S. M., Horsley, V., and King, M. C. (2015). Nuclear-cytoskeletal linkages facilitate cross talk between the nucleus and intercellular adhesions. J. Cell Biol. 209, 403–418.

Stewart-Hutchinson, P. J., Hale, C. M., Wirtz, D., and Hodzic, D. (2008). Structural requirements for the assembly of LINC complexes and their function in cellular mechanical stiffness. Exp. Cell Res. 314, 1892–1905.

Stroud, M. J., Banerjee, I., Veevers, J., and Chen, J. (2014). Linker of nucleoskeleton and cytoskeleton complex proteins in cardiac structure, function, and disease. Circ. Res. 114, 538–548.

Su, Y., Xia, W., Li, J., Walz, T., Humphries, M. J., Vestweber, D., Cabañas, C., Lu, C., and Springer, T. A. (2016). Relating conformation to function in integrin α5β1. Proc Natl Acad Sci U S A 113, E3872–E3881.

Taegtmeyer, H., Sen, S., and Vela, D. (2010). Return to the fetal gene program: a suggested metabolic link to gene expression in the heart. Ann. N. Y. Acad. Sci. 1188, 191–198.

Thakar, K., May, C. K., Rogers, A., and Carroll, C. W. (2017). Opposing roles for distinct LINC complexes in regulation of the small GTPase RhoA. Mol. Biol. Cell 28, 182–191.

Travers, J. G., Kamal, F. A., Robbins, J., Yutzey, K. E., and Blaxall, B. C. (2016). Cardiac Fibrosis: The Fibroblast Awakens. Circ. Res. 118, 1021–1040.

Varney, S. D., Betts, C. B., Zheng, R., Wu, L., Hinz, B., Zhou, J., and Van De Water, L. (2016). Hic-5 is required for myofibroblast differentiation by regulating mechanically dependent MRTF-A nuclear accumulation. J. Cell. Sci. 129, 774–787.

Wang, Q., Lin, J. L.-C., Wu, K.-H., Wang, D.-Z., Reiter, R. S., Sinn, H. W., Lin, C.-I., and Lin, C.-J. J. (2012). Xin proteins and intercalated disc maturation, signaling and diseases. Front Biosci (Landmark Ed) 17, 2566–2593.

Wilsbacher, L. D., and Coughlin, S. R. (2015). Analysis of cardiomyocyte development using immunofluorescence in embryonic mouse heart. J Vis Exp, e52644–e52644.

Wu, W., Muchir, A., Shan, J., Bonne, G., and Worman, H. J. (2011). Mitogen-activated protein kinase inhibitors improve heart function and prevent fibrosis in cardiomyopathy caused by mutation in lamin A/C gene. Circulation 123, 53–61.

Young, J. L., Kretchmer, K., Ondeck, M. G., Zambon, A. C., and Engler, A. J. (2014). Mechanosensitive kinases regulate stiffness-induced cardiomyocyte maturation. Sci Rep 4, 6425.

Zhou, C. et al. (2017). Novel nesprin-1 mutations associated with dilated cardiomyopathy cause nuclear envelope disruption and defects in myogenesis. Hum. Mol. Genet. 26, 2258–2276.

Zwerger, M., Jaalouk, D. E., Lombardi, M. L., Isermann, P., Mauermann, M., Dialynas, G., Herrmann, H., Wallrath, L. L., and Lammerding, J. (2013). Myopathic lamin mutations impair nuclear stability in cells and tissue and disrupt nucleo-cytoskeletal coupling. Hum. Mol. Genet. 22, 2335–2349.

