## Supplemental Figures 1-3 for "Ablation of SUN2-containing LINC complexes drives cardiac hypertrophy without interstitial fibrosis"

**Stewart et al. Supplemental Figures 1-3**

**Supplemental Figure 1. *Sun2-/-* mice lack SUN2 in the myocardium without compensation by SUN1**

**(A)** Frozen P50 WT and *Sun2-/-* cardiac left ventricle tissue was sectioned and stained with antibodies against SUN2 (grey) and counterstained with Hoechst 33342 (blue). SUN2 is present at nuclear envelopes in WT tissue (red arrow), but absent from *Sun2-/-* tissue. **(B)** qPCR analysis of *Sun1* mRNA levels from 13-month-old left ventricular total RNA, indicating that SUN1 is present in both WT and *Sun2-/-* ventricle tissue. WT and *Sun2-/-* Ct values were normalized to GAPDH. Fold change of *Sun2-/-* mRNA levels was determined by calculating the $2^{\Delta\Delta Ct}$ after taking the mean of WT ΔCt values from 5 mice. n = 4 *Sun2-/-* mice. Statistical significance determined by unpaired, two-tailed *t* test and found to not be significant.

**Supplemental Figure 2. *Sun2-/-* myocardium exhibits cardiomyocyte adhesion defects**

**(A)** Transmission electron micrographs (TEM) of P50 WT and *Sun2-/-* left ventricle tissue. Note the reduced electron density along the intercalated disc membrane in *Sun2-/-* compared to WT tissue (upper panels, red arrows; lower panels, red arrowheads), as well as regions lacking myofilament association with the intercalated disc in *Sun2-/-* tissue (lower panels, arrows). **(B and C)** Frozen P50 WT and *Sun2-/-* cardiac left ventricle tissue was sectioned and stained with antibodies against desmoplakin I/II (DPI/II) **(B)**, β-catenin **(C)**, and counter-stained with wheat germ agglutinin (WGA, red). Upper panels display 100x magnification views of the tissue; lower panels are insets of the white-boxed regions. Both DPI/II and β-catenin exhibit abnormally expanded, striated localization at intercalated disc sites in *Sun2-/-* tissue (arrowheads). **(D)** Frozen P13 WT and *Sun2-/-* ventricle tissue was stained with antibodies against DPI/II, revealing similar patterns of staining in both genotypes. **(E)** Frozen P50 WT and *Sun2-/-* cardiac left ventricle tissue was sectioned and stained with antibodies against total β1-integrin (upper panels) or against the ligand-bound, active β1-integrin 9EG7 conformational epitope (lower panels). Images were pseudo-colored as in Figure 4A. Images are representative of 2 additional biological repeats of the staining shown in Figure 4A. Note the increased intensity of active β1-integrin staining in the *Sun2-/-* tissue. **(F)** Frozen P13/P14 WT and *Sun2-/-* cardiac left ventricle tissue was sectioned and stained with 9EG7 antibodies against active β1-integrin, revealing similar levels in both genotypes. Images are representative of 2 biological repeats. Images were pseudo-colored as in Figure 4A. **(G)** Frozen 13 month-old WT and *Sun2-/-* cardiac left ventricle tissue was sectioned and stained with 9EG7 antibodies against active β1-integrin, revealing similar levels in both genotypes. Images are representative of 3 biological repeats. Images are pseudo-colored as in Figure 4A.

**Supplemental Figure 3. *Sun2-/-* mice exhibit elevated AKT/MAPK signaling**

(A-D) Full immunoblots corresponding to Figure 4B-E. Ponceau staining of total protein revealed even loading of samples. Representative blots are shown. (A) Ventricular tissue was isolated from 5 WT and 4 *Sun2-/-* 13 month-old mice and total protein lysates were obtained. SDS-PAGE and immunoblotting with antibodies against phosphorylated AKT (pAKT) or AKT revealed similar levels of pAKT in *Sun2-/-* tissue. We note that some *Sun2-/-* mice exhibited higher levels of pAKT/AKT ratios than WT, but this effect was not consistent across all mice examined. (B) Ventricular lysates from 13 month-old mice as in (A) were immunoblotted with antibodies against phosphorylated S6 (pS6) and S6, revealing elevated levels of pS6 in *Sun2-/-* mice. Full blots from Figure 4C as indicated. (C) Ventricular tissue was isolated from 3 WT and 3 *Sun2-/-* P50 mice and total protein lysates were obtained. SDS-PAGE and immunoblotting with antibodies against pS6, S6, pAKT, or AKT revealed elevated levels of pAKT but similar levels of pS6 in *Sun2-/-* tissue. Upper panels represent longer exposures of the same blots shown in the lower panels. Full blots from Figure 4B as indicated. (D) Ventricular lysates from P50 and 13 month-old mice as in (A,B) were immunoblotted with antibodies against phosphorylated ERK1/2 (pERK1/2) or ERK2, revealing elevated levels of pERK1/2 in P50 *Sun2-/-* tissue but similar levels of pERK1/2 in 13 month-old *Sun2-/-* tissue compared to WT.

#### Supplemental Figure 1.

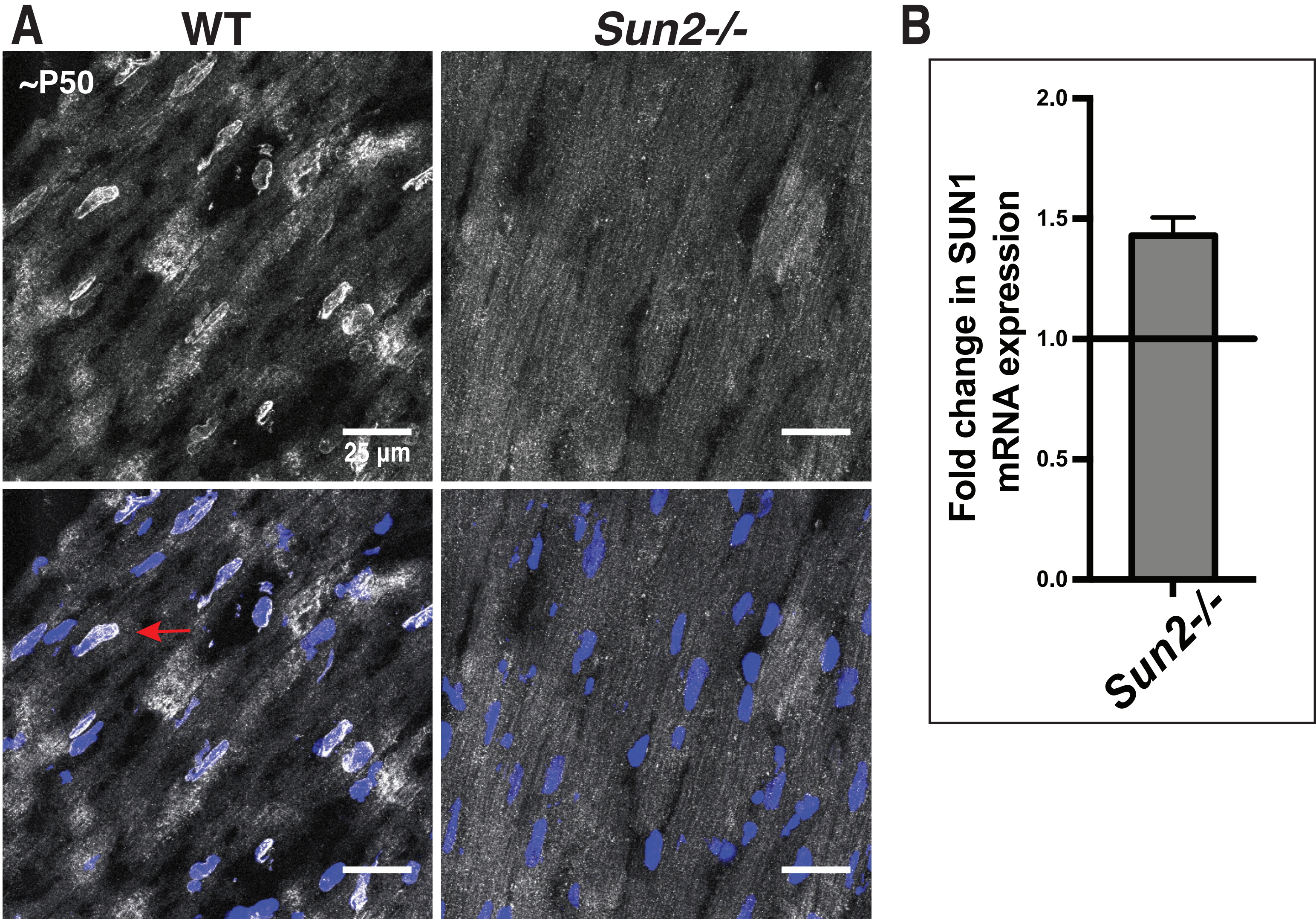

### Supplemental Figure 2.

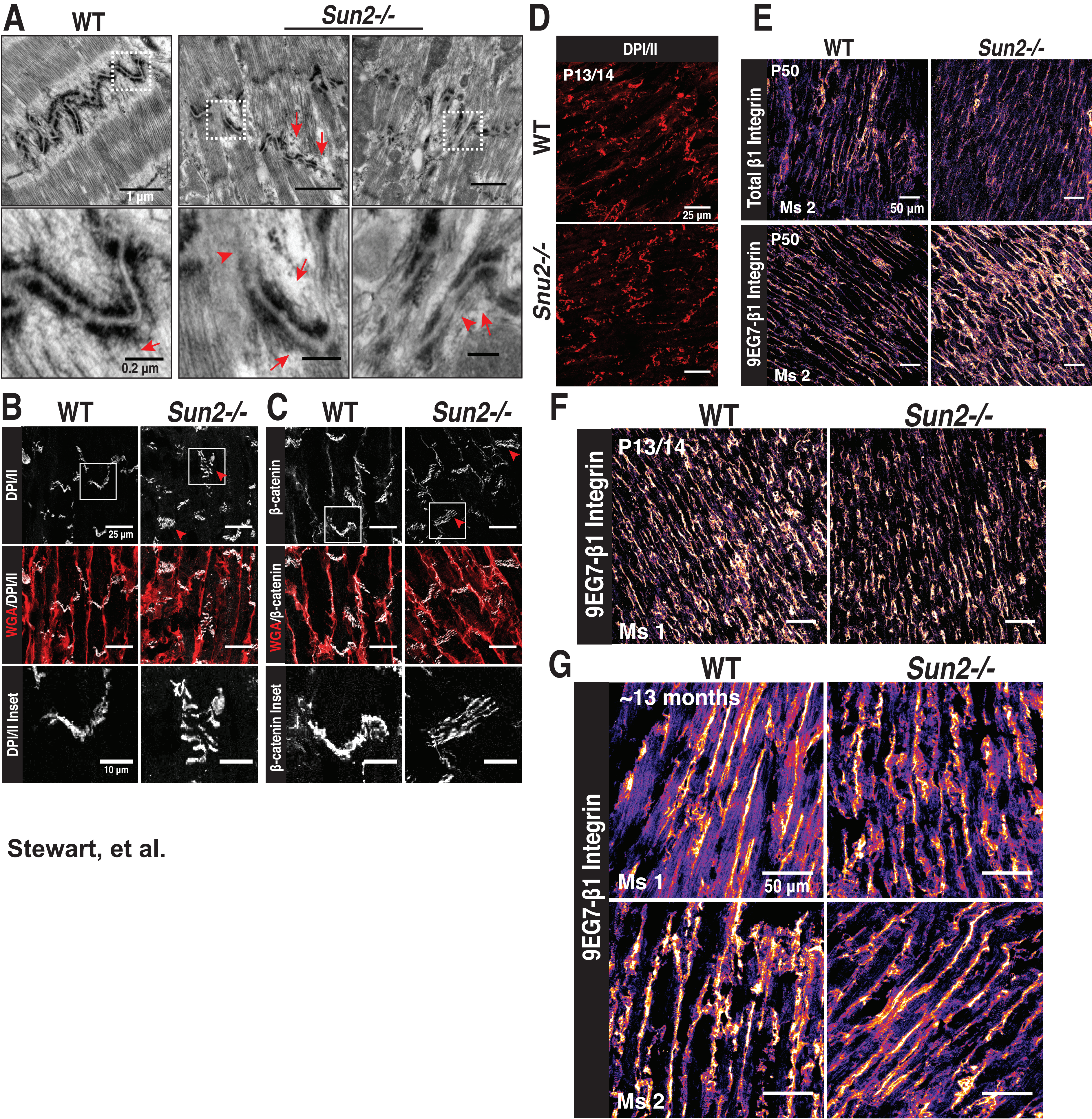


Stewart, et al.

Supplemental Figure 3.

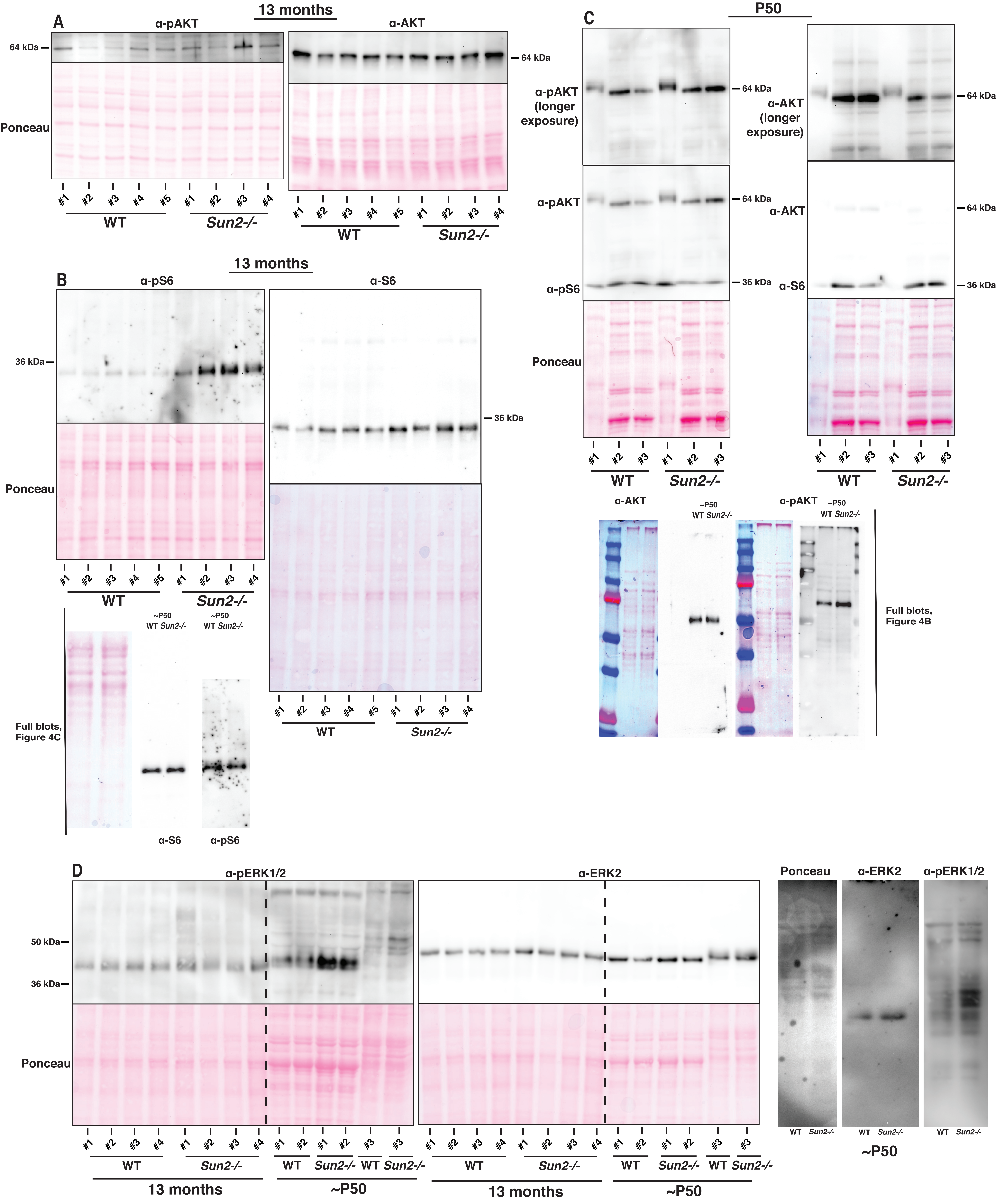
